# Swimming, fast and slow: strategy and survival of bacterial predators in response to chemical cues

**DOI:** 10.1101/2020.11.11.377200

**Authors:** M Carlson, S L Seyler, S Pressé

## Abstract

*Bdellovibrio bacteriovorus* is a predatory bacterium that preys upon gram-negative bacteria. As such, *B. bacteriovorus* has the potential to control antibiotic-resistant pathogens and biofilm populations. To survive and reproduce, *B. bacteriovorus* must locate and infect a host cell. However, in the temporary absence of prey, it is largely unknown how *B. bacteriovorus* modulate their motility patterns in response to physical or chemical environmental cues to optimize their energy expenditure. To investigate *B. bacteriovorus’* predation strategy, we track and quantify their motion by measuring speed distributions and velocity autocorrelations as a function of starvation time. An initial unimodal speed distribution, relaxing to that expected for pure diffusion at long times, may be expected. Instead, we observe a complex, non-Brownian, search strategy as evidenced by distinctly bimodal speed distributions. That is, for an increasing amount of time over which *B. bacteriovorus* is starved, we observe a progressive re-weighting from a fast mode to a slow mode in the speed distribution obtained over consecutive frames. By contrast to its predator, *B. bacteriovorus’* prey, *Escherichia coli* exhibits almost immediate decrease to a speed expected from passive diffusion following resuspension from rich to poor media. Distributions of trajectory-averaged speeds for *B. bacteriovorus* are largely unimodal, indicating nontrivial *switching* between fast and slow swimming modes within individual observed trajectories rather than there being distinct fast and slow populations. We also find that *B. bacteriovorus’* slow speed mode is not merely caused by the diffusion of inviable bacteria as subsequent spiking experiments show that bacteria can be resuscitated and bimodality restored. Indeed, starved *B. bacteriovorus* may modulate the frequency and duration of active swimming as a means of balancing energy consumption and procurement. Our results are evidence of a nontrivial predation strategy, which contrasts with the comparatively simple search pattern of its prey, in response to environmental cues.

**SIGNIFICANCE:** *Bdellovibrio bacteriovorus* is a predatory bacterium that is poised to help control gram-negative bacterial populations in environmental and clinical settings. In order to locate its prey in solution, *B. bacteriovorus* must expend energy in order to fight hydrodynamic drag. This raises the question as to how *B. bacteriovorus* should expend its energy reserves in the absence of chemical cues from its prey. Here, we show that *B. bacteriovorus* adapts its motility to minimize energy expenditure (due to fighting drag in swimming) upon prolonged starvation by exploiting two modes of motility. This is in sharp contrast to its prey, *E. coli*, which shows little active motility under starvation conditions.

## INTRODUCTION

Bacterial cues drawn from the environment—mediated through a number of factors such as hydrodynamic interactions (1, 2), external flows (3–5) or other direct chemical signaling and quorum sensing (6–12)—are critical in understanding cell-cell interaction and emergent collective behaviors of biomedical interest (7, 13, 14) such as biofilm formation (15) and bacterial swarming (16, 17).

Here our focus is in understanding the hunting strategy of predatory bacteria, such as *Bdellovibrio* and like organisms (BALOs), driven by external cues. Understanding such strategies is helpful downstream in exploiting BALOs for a number of tasks including the degradation of hazardous microbial biofilms on surfaces (15, 18–22), purifying water and soil (23–25), and serving as a “living antibiotic” (14, 26, 27) by reducing pulmonary bacterial infections in rats (28), gut infections in poultry (29), corneal infections in rabbit (30) and infections in other animal models such as zebra fish (31). Indeed, the bacterial predator, *Bdellovibrio bacteriovorus*, has been shown to significantly decrease populations of many other species of gram-negative bacteria (32, 33) across habitats (14, 15, 18, 19, 23–25, 29–31, 34, 35).

The biphasic life cycle of *B. bacteriovorus* involves an attack phase in which predatory *B. bacteriovorus* locate their bacterial prey and enter the periplasm where they complete the replicative growth phase, ending with the progeny lysing from the host (36, 37, 37–42). Although *B. bacteriovorus* was discovered half a century ago (43–45) and is now studied as a model predator, it is largely unknown how *B. bacteriovorus* locate individual prey. For instance, previous studies suggested that *B. bacteriovorus* bumps into its prey at random (19, 23, 46–51).

In previous work we re-visited this hypothesis (2) and investigated the role hydrodynamics plays on *B. bacteriovorus’* hunting strategy. Our own prior efforts built upon recent literature suggesting that bacteria sense and respond to environmental hydrodynamic flows (3–5) and self-generated flows (1, 52–57). In particular we found that, on account of *B. bacteriovorus’* unusually high speed, passive hydrodynamic forces could drive it towards surfaces where prey tend to be present in larger numbers (2), thereby improving the odds of a chance collision.

This finding raises the following important question: given that hydrodynamic forces induced by *B. bacteriovorus’* unusually high speeds are critical in bringing it toward prey, how might *B. bacteriovorus*, in the absence of prey in its environment, allocate its energy reserves to overcome hydrodynamic drag as it propels itself through solution in search of prey? A naive search strategy where active motility persists until the available energy—used to maintain metabolic homeostasis, as well as overcoming both translational and rotational hydrodynamic drag—is expended (12) seems patently suboptimal. Such a scenario suggests that *B. bacteriovorus would not be revivable* once energy resources are depleted and, possibly, that its proteome might be cannibalized in an effort to locate prey by active swimming (58). On the other hand, a more sophisticated use of energy would involve the active modulation of motility as a function of both internal energy reserves (i.e., age) and environmental cues (i.e., potential sources of energy and nutrition) prior to reaching a point of no return. That is, the point at which *B. bacteriovorus* can no longer fight hydrodynamic drag in search of prey despite external cues suggesting the availability of prey.

Here we address this question by monitoring changes in large numbers of *B. bacteriovorus* trajectories under starvation conditions as a function of age and buffer conditions. We found that key dynamical features—instantaneous and trajectory-averaged speed distributions and autocorrelation functions—monitored over the course of forty hours suggest behavior consistent with an exploitative search strategy. That is, a complex, non-Brownian, search pattern that is modulated over time and evidenced by the slower decay of the speed autocorrelation recovered after extended periods of starvation by *B. bacteriovorus*.

We corroborated these results by subsequently performing spiking experiments in which starved bacteria are resuspended in rich media, showing that *B. bacteriovorus* can be resuscitated and their active motility can be restored. A key aspect of our work is the comparison of *B. bacteriovorus’* motility pattern under starvation to its prey *E. coli*. Our results help shed light on the difficult compromises that *B. bacteriovorus* must make when hunting in the absence of environmental cues and how quantitative speed modulation can be used to make the most of unfavorable environmental conditions.

## MATERIALS AND METHODS

### Bacterial strains and growth conditions

In order to avoid any possible variations in the motile behavior of *B. bacteriovorus* due to expression of a fluorescent protein, the wild type strain 109 (BV Stolp and Starr, ATTC No. 15143; American Type Culture Collection, Manassas, VA) was utilized to carry out these studies. *Escherichia coli* strain OP50 which was grown in LB (10 g/L yeast extract, 20 g/L tryptone, 20 g/L sodium chloride, pH 7.5) at 37°C on a 300-rpm shaker was used as the prey of *B. bacteriovorus*. After multiple washes with HEPES medium, the OD (600 nm) was taken of the *E. coli* and the bacteria were stored at 4°C until use for synchronization cultures as described below.

### Synchronization of *Bdellovibrio bacteriovorus*

Modifications to the synchronization protocols outlined in the literature (59, 60) allow for the synchronized observation of many *B. bacteriovorus* of the same age. This was achieved by growing *B. bacteriovorus* in nutrient-poor medium (HM) (25 mM HEPES, pH 7.4, 3 mM CaCl_2_, 2 mM MgCl_2_) containing approximately 10^8^ (OD 1) *E. coli* cells overnight at 28°C on a 300-rpm shaker. The next day, *B. bacteriovorus* was isolated by centrifugation at 7900 rpm for 30 minutes at 4°C. The predators were then resuspended in fresh HM buffer containing *E. coli* cells at a 0.7 OD. An hour after introduction to fresh *E. coli*, the formed bdelloplasts were isolated and resuspended in fresh media several times via centrifugation (2000 rpm for 10 minutes at room temperature). This eliminated the majority of leftover predators. After four additional hours, the lysed progeny were collected by passing the culture through a 0.45 µm Millex HP syringe filter twice. The isolated *B. bacteriovorus* remained in nutrient-poor media to begin starvation conditions.

The effectiveness of the isolation method was tested by passing an *E. coli* OP50 culture of OD 1 through the filter and adding nutrient-rich media. The culture was monitored for several days, no bacteria grew, showing that *E. coli* was effectively removed by filtration. In addition, an OP50 culture was starved in HM buffer and then spiked as the *B. bacteriovorus* were in the runs described below. The speed distributions after spiking were compared to that of *B. bacteriovorus* and can be seen in Fig. S22 and Fig. S23. All *E. coli* OP50 are diffusive unlike the predator *B. bacteriovorus*.

### Starvation of *B. bacteriovorus* experiments

The synchronized *B. bacteriovorus* was obtained as described above. The culture remained shaking at 28°C on a 300-rpm shaker during the total duration of the experiment (approximately 48 hours). A small aliquot was taken from the culture every hour for the sample preparation and data acquisition described below.

### Spiking of *B. bacteriovorus* with rich media experiments

The *B. bacteriovorus* was synchronized as described above. Four hours after the culture was filtered, the culture was divided into three 2 mL total volumes: a control, and two cultures that were spiked with rich media. The *B. bacteriovorus* is spiked with nutrient-rich media, LB, (10 g/L yeast extract, 20 g/L tryptone, 20 g/L NaCl, pH 7.5) at Hour 4 and Hour 20 after filtration. The cultures that are spiked have 1.75 mL of *B. bacteriovorus* added and 0.25 mL of rich media. The cultures remained shaking at 28°C on a 300-rpm shaker during the total duration of the experiment. Small aliquots were taken from the three cultures for measurements.

### Memory of *B. bacteriovorus* experiments

The *B. bacteriovorus* was synchronized according to the “synchronization of *B. bacteriovorus*” section. Six hours after filtration, the culture was divided into three 2 mL total volumes: a control, and two cultures that were spiked with rich media at Hour 6. The cultures that are spiked have 1.75 mL of *B. bacteriovorus* added and 0.25 mL of rich media. The cultures are observed 15 minutes after exposure. To initiate starvation conditions again, all three cultures were centrifuged (30 minutes after initial exposure time) at 13,000 rpm for 4 minutes at room temperature. The control culture and one of the spiked cultures were resuspended in 2 mL fresh HM. The other spiked culture was resuspended in 1.75 mL of HM and 0.25 mL of LB. (61). The cultures remained shaking at 28°C on a 300-rpm shaker during the total duration of the experiment. Small aliquots were taken from the three cultures for measurements. The effects of centrifugation can be seen in Fig. S18.

### Starvation of *E. coli* experiments

*E. coli* strain MG1655 was used for the starvation experiments which has been used in previous studies (2, 33). A 5 mL culture was grown overnight from a single colony. The next day, the OD was measured (approximately OD 3) and 20 µL was added for a 6 mL total volume culture containing LB. After 2 hours (OD 0.1), the *E. coli* was seen swimming and videos of their motion was acquired (Hour -1). To initiate starvation conditions, the culture was centrifuged at 2000 rpm for 10 minutes at room temperature and resuspended in HM. A separate control culture was resuspended in LB to compare the effect of centrifugation. Due to the rod-shape of *E. coli*, they were observed at the coverslip for better tracking. Similar behavior after initiation of starvation was seen in the midplane.

### Sample preparation

A small volume (10 µL) of the culture was aliquoted onto a microscope cover slip containing vacuum grease along the outer edge and secured with a slide. Focusing on the midplane (approxiamtely 70 µm from the surface) allowed for the observation of the bacteria without restricting their movement due to a confined vertical depth (2).

### Data acquisition

Using an inverted phase contrast microscope (Nikon, Melville, NY), the bacteria were imaged with a 1024×1024 pixel ROI using a 60X (1.4 NA) oil immersion objective. Each hour data was collected by recording six twenty-second videos at 8 ms exposure using a Hamamatsu ORCA-Flash4.0 3V sCMOS camera. These videos were then processed to obtain velocity distributions over thousands of bacterial trajectories as described in the “data analysis” section.

## Data analysis

The motility of the bacteria was tracked by utilizing a package for feature detection called Trackpy (62), which provides computational tools to locate features (e.g., bacteria) in raw image data and then link those features into trajectories. (A trajectory is a time-sequence of changes, from frame to frame, in the spatial location of a feature in the focal plane, i.e., a single bacterium.) Two of the main parameters for locating features are the diameter and minmass. The diameter is determined based on the size of the object tracked and the minmass is the minimum integrated brightness used to eliminate spurious features. As *B. bacteriovorus* is approximately 1 micron (i.e., 11 pixels), we chose an all inclusive 15 pixel diameter. The minmass varies between runs given the overall brightness of the video. Changes in this value may affect the overall number of bacteria tracked within the focal plane, but would not influence the overall motile behavior. To link the bacteria, the search range was always set to 20 as the bacteria should not travel further than this number of pixels between consecutive frames (19). Lastly, in filtering trajectories, we ignored any traces that were shorter than 10 consecutive frames in order to get rid of any spurious trajectories. As bacteria move in and out of focus, there is a sampling bias in speed measurements at the start and end of trajectories towards slow speeds (i.e., a bias towards a larger velocity component *orthogonal* to the focal plane). Removing the first and last two frames of all trajectories was sufficient to mitigate this bias (Fig. S19, Table S1).

Window averaging of the bacterium’s position was calculated in order to more accurately track its movement. The effect of different windows can be seen in Fig. S6. Based on these results, a window of three was utilized for all located bacteria. As windowing essentially decreases the number of original data points, we also tested down-sampling (decimating) the data to conclude that the shape of the distributions did not greatly vary (Fig. S21).

Once each trajectory is recorded as a time series of 2-dimensional coordinates, the distance the feature moves between each consecutive frame and the exposure time of the camera was used to calculate the instantaneous speeds. A central difference of the positions with respect to time was used to get a more accurate estimate of these speeds. The average speed for each trajectory was obtained by taking the average of the instantaneous speeds of each bacterium’s trajectory. The VACF was also calculated using the *xy*-components of the velocity to gain insight into the underlying dynamics of *B. bacteriovorus’* motility, information which speed distributions alone cannot provide (see Eq. (4) for details).

## RESULTS AND DISCUSSION

To see how *B. bacteriovorus* adapts to environmental cues, we first observed the swimming behavior of synchronized *B. bacteriovorus* in the absence of prey and nutrients. Under such starvation conditions, we monitored the instantaneous speeds between two consecutive frames and the average speed over each bacterium’s trajectory (Fig. 1). We uncover a reproducible bimodal instantaneous speed distribution with peaks at roughly 10 µm s^−1^ and 40 µm s^−1^ with the relative weight of each peak being sensitive to *B. bacteriovorus’* age and buffer conditions. We calculated the instantaneous velocity autocorrelation function (VACF) to gain quantitative insight into whether the bacteria exhibit a change in directed motility as a function of *B. bacteriovorus’* age (Fig. 3). Or, put differently, at what age *B. bacteriovorus* exhibits a rapid decay in its VACF anticipated for a purely Brownian particle. We also calculate the power dissipation to further determine how *B. bacteriovorus* budgets its limited energy resources (Fig. 4). This allows us to gain insight into how these predators adapt their motility patterns in the absence of external stimuli. These results are contained in the “starvation of *B. bacteriovorus*” section.

**Fig 1.**
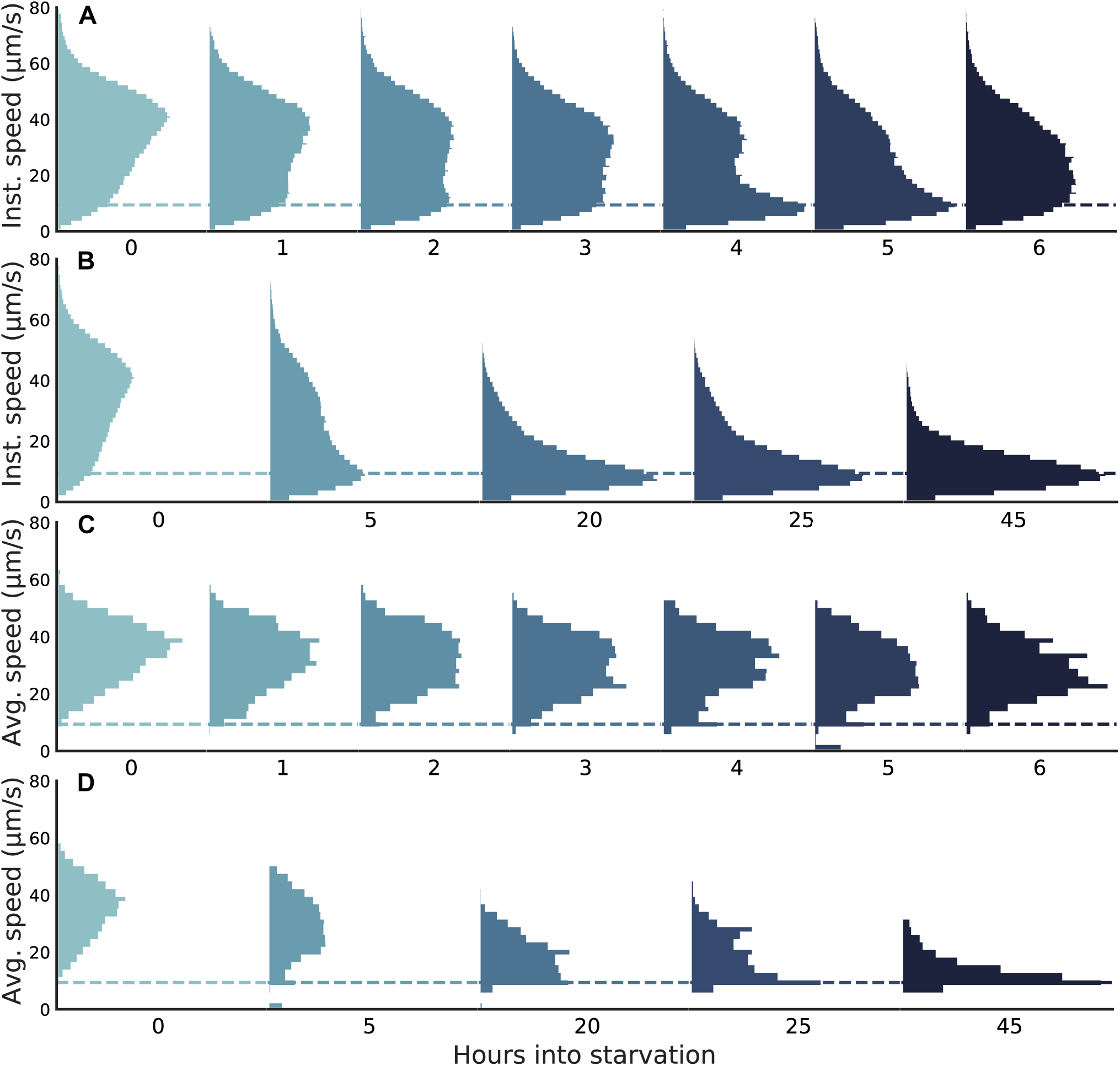
Under starvation conditions, the instantaneous and average speed distributions of *B. bacteriovorus* shift across time. (A) Instantaneous speed distributions are bimodal with a substantial shift toward lower-speed observations over the first 6 hours. (B) After up to 45 hours of starvation, the instantaneous speed distribution relaxes to a unimodal distribution about the expected diffusive speed for a Brownian particle of *B. bacteriovorus’* size (≈ 9.40 µm s^−1^, dashed lines). (C) Distributions of speeds averaged over individual trajectories are largely unimodal over the first 6 hours. (D) Average speed distributions over 45 hours eventually relax to the expected diffusive speed. The sampling statistics for each histogram can be found in S3 Text.

In the subsequent section, we introduced *B. bacteriovorus* to nutrient-rich media [lysogeny broth, LB]—effectively providing them with an external energy source—in order to determine whether the bacteria were merely inviable or strategically swimming more slowly at later times. We monitored the changes in the speed distributions (Fig. 5) and the VACF Fig. 6) as compared to a starving control of equal age. To quantify the energy expenditure for motility, we computed the average power dissipation as a function of age, as well as in the presence or absence of LB (Fig. 7).

To further investigate the nature of *B. bacteriovorus’* search strategy (i.e., the bimodal behavior), we monitored *B. bacteriovorus* ’ behavior with a cycle of starvation, rich media exposure, and then re-starvation. This resulted in the bacteria re-populating the fast speed and then the slow speed mode, respectively (Fig. 8).

To compare the behavioral changes of a predator versus prey under starvation conditions, we monitored the motility of *E. coli* in nutrient-poor media (Fig. 9). Our conclusions are consistent with the expectation that *E. coli* can slow down as it is not required to expend energy to find prey (63–65).

### Starvation of *B. bacteriovorus*

In Fig. 1, we show the instantaneous and average speed distributions as a function of age. As described in the Methods, Hour 0 is observed ten minutes after filtering. Starting from Hour 0, the instantaneous speeds exhibit a bimodal distribution, with peaks around 40 µm s^−1^ and 10 µm s^−1^. The latter peak is centered around the expected diffusive speed of a spherical particle the size of *B. bacteriovorus*, as predicted by the Stokes-Einstein relation (Fig. 1, dashed line) (see S1 Text for details).

As bacteria age over the first six hours under starvation conditions, we observe a decrease in the higher-speed population and an increase in the lower-speed population of our bimodal distribution. The observed shifts suggest the possibility that the bacteria are actively adjusting their swim speed in response to persisting starvation conditions. The evolution of the speed distribution, from Hour 0 up to Hour 45, is shown in Fig. 1B (with bimodality shifting to unimodality around the 20 Hour mark). In the methods, we describe tracking details.

To determine whether there are two bacterial populations or two modes of swimming for each bacterium contributing to the bimodal instantaneous distributions (Fig. 1A,B), the average velocity over each bacterium’s trajectory was calculated (Fig. 1C).

The speed distributions have a similar average velocity over the first six hours. This indicates that the speed of each bacterium is more similar to that of other bacteria than the speeds sampled within its own trajectory. However, just as before, the average velocity approaches the diffusive speed after Hour 20 (Fig. 1D, dashed line), suggesting that the bacteria are passively diffusing. The absence of a well defined unimodal speed distribution is corroborated by direct visual inspection of bacterial trajectories; representative traces, exhibiting both fast and slow speeds within individual trajectories, are shown in Fig. 2. Additional runs can be found in the supplementary information (Fig. S1, Fig. S2, and Fig. S3).

**Fig 2.**
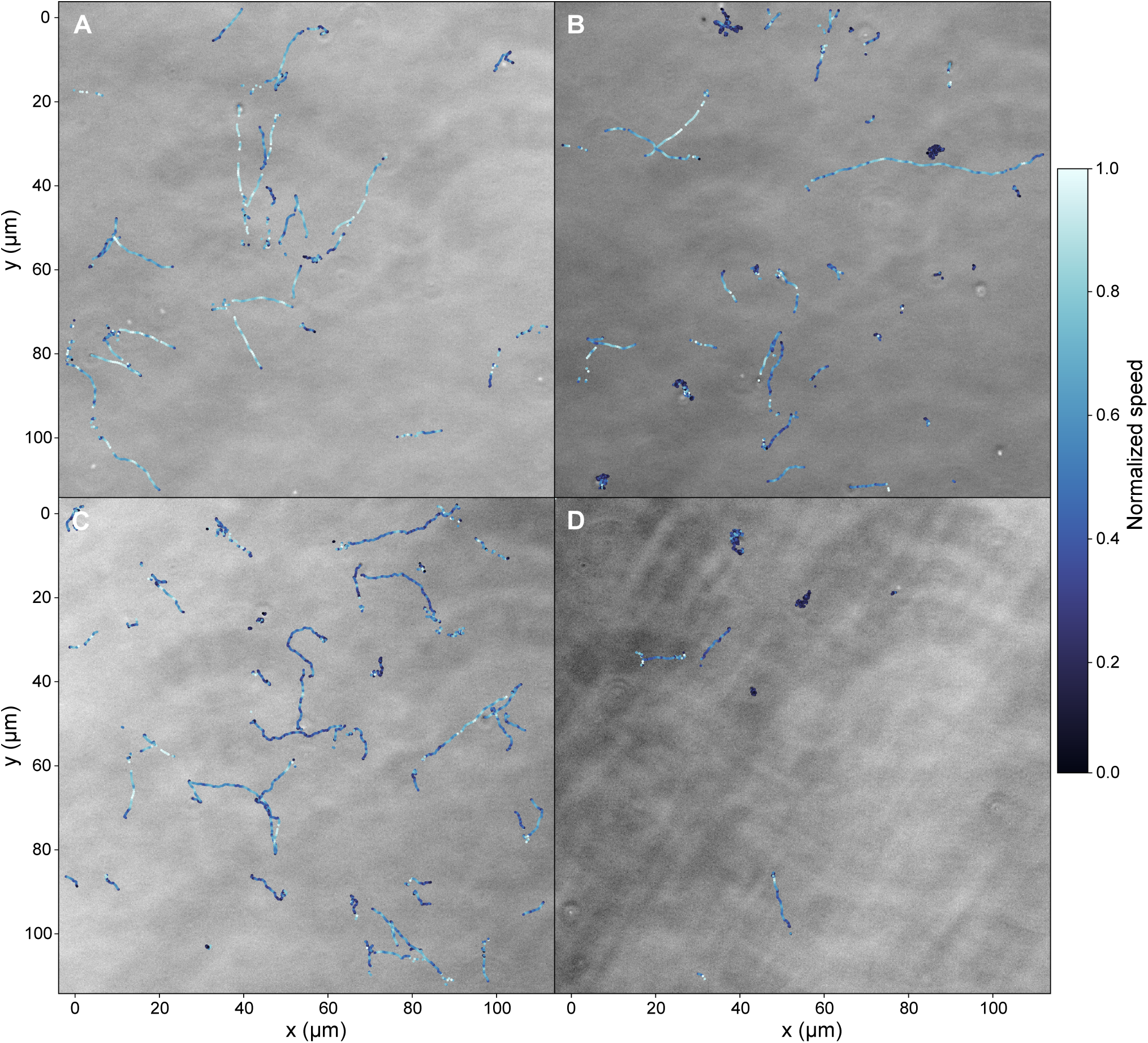
Representative *B. bacteriovorus* trajectories depicting instantaneous velocities. Bacterial trajectories are obtained for the first 500 frames at Hour 0 (A), Hour 4 (B), Hour 6 (C), and Hour 20 (D). The colors within each trajectory correspond to the normalized instantaneous speed.

Under starvation conditions, we were still left to wonder whether, after the first 6 hours (Fig. 1), *B. bacteriovorus* are inviable or passively diffusing but still viable. In order to determine which of these two scenarios is realized, we calculate the instantaneous VACF; see Fig. 3.

**Fig 3.**
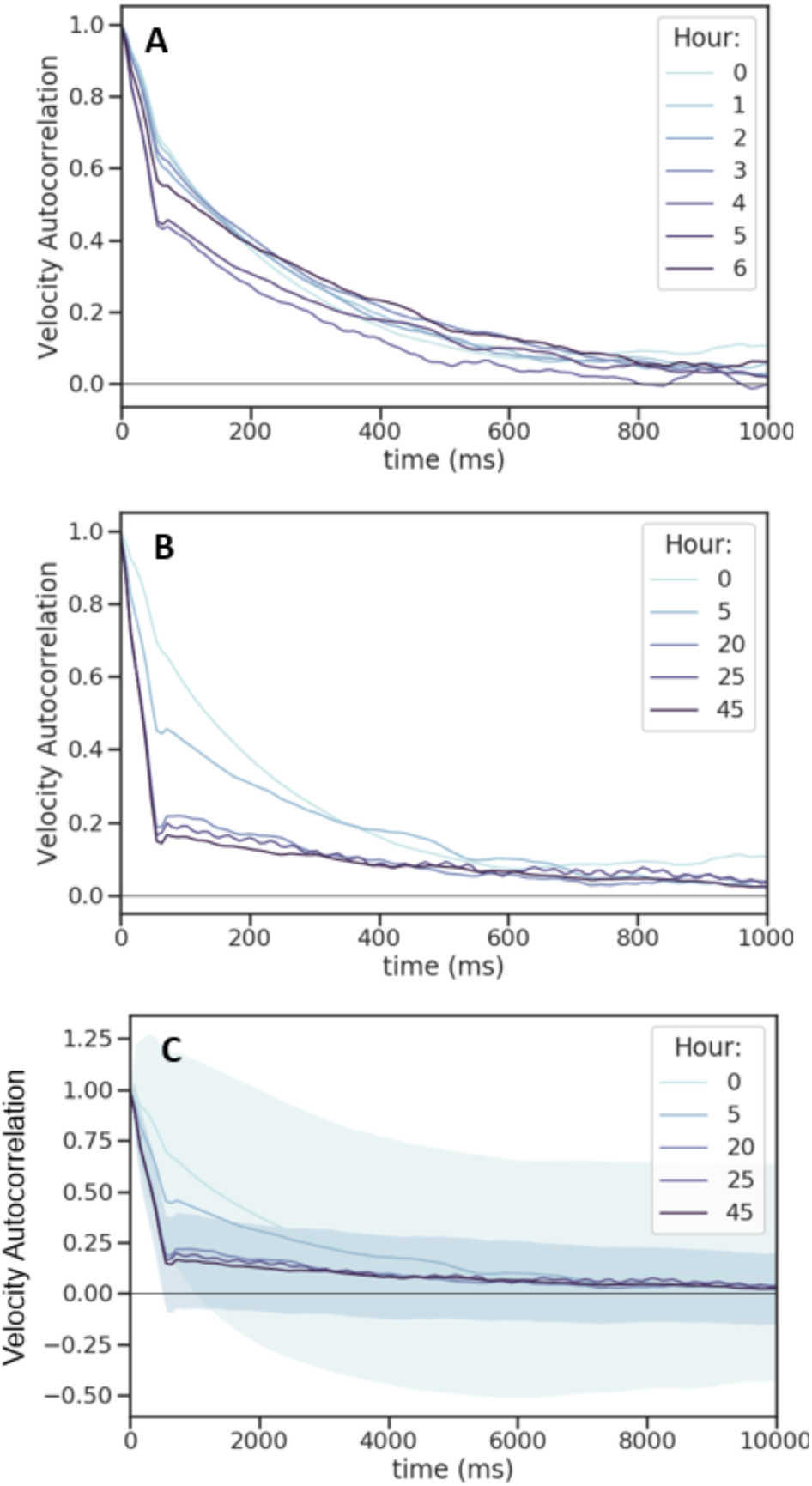
Under starvation conditions, *B. bacteriovorus’* velocity decorrelates increasingly rapidly as it ages over 45 hours of observation. (A) From Hour 0 to Hour 6, the decay of the instantaneous velocity time autocorrelation over hundreds of milliseconds indicate that the bacteria are actively self-propelling. (B) As the bacteria age from Hour 0 to Hour 45, their velocities are largely decorrelated on timescales of tens of milliseconds. (C) The results of panel (B) are depicted with error bands about hour 0 and hour 45. Details of VACF calculations are provided in Eq. 1 of S2 Text. The sampling statistics for each histogram can be found in S3 Text.

We find that the VACF decays more rapidly as *B. bacteriovorus* ages (Fig. 3A). This is consistent with the notion that the motile behavior shifts toward (passive) thermal diffusion. In particular, the decay from Hour 0 to Hour 6 (Fig. 3B) is gradual as compared to the rapid decay at Hour 20 and later. When observing the speed distributions of *B. bacteriovorus* at Hour 20 and Hour 40, we notice negligible differences. Yet, when comparing the VACF at both of these hours, we see a slight indication of correlation at Hour 20 by contrast to minimal correlation by Hour 40 (Fig. 3C). These results (in addition to separate runs Fig. S4, Fig. S5, and Fig. S6), taken together, suggest that *B. bacteriovorus* is not only be revivable after many hours of starvation, but can actively alter its motile behavior in response to environmental cues. This will be explored in the subsequent section.

Before we do so, we first comment on how *B. bacteriovorus* budgets its limited energy resources for active swimming by calculating the Stokes power dissipation (12),

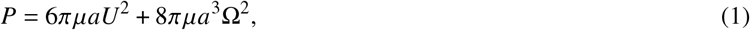

where *µ* is the dynamic viscosity of the fluid, *a* the (bacterial) radius, and *U* and Ω are the forward and angular velocities, respectively. For simplicity, we make the usual assumption that a bacterium’s angular velocity is proportional to the rotational speed of its flagellum (55, 66), which is furthermore proportional to its forward speed (55, 66). Under these assumptions, Eq Eq. (1) predicts that the instantaneous power dissipation grows quadratically with velocity *U*, the results of which are shown in Fig. 4.

**Fig 4.**
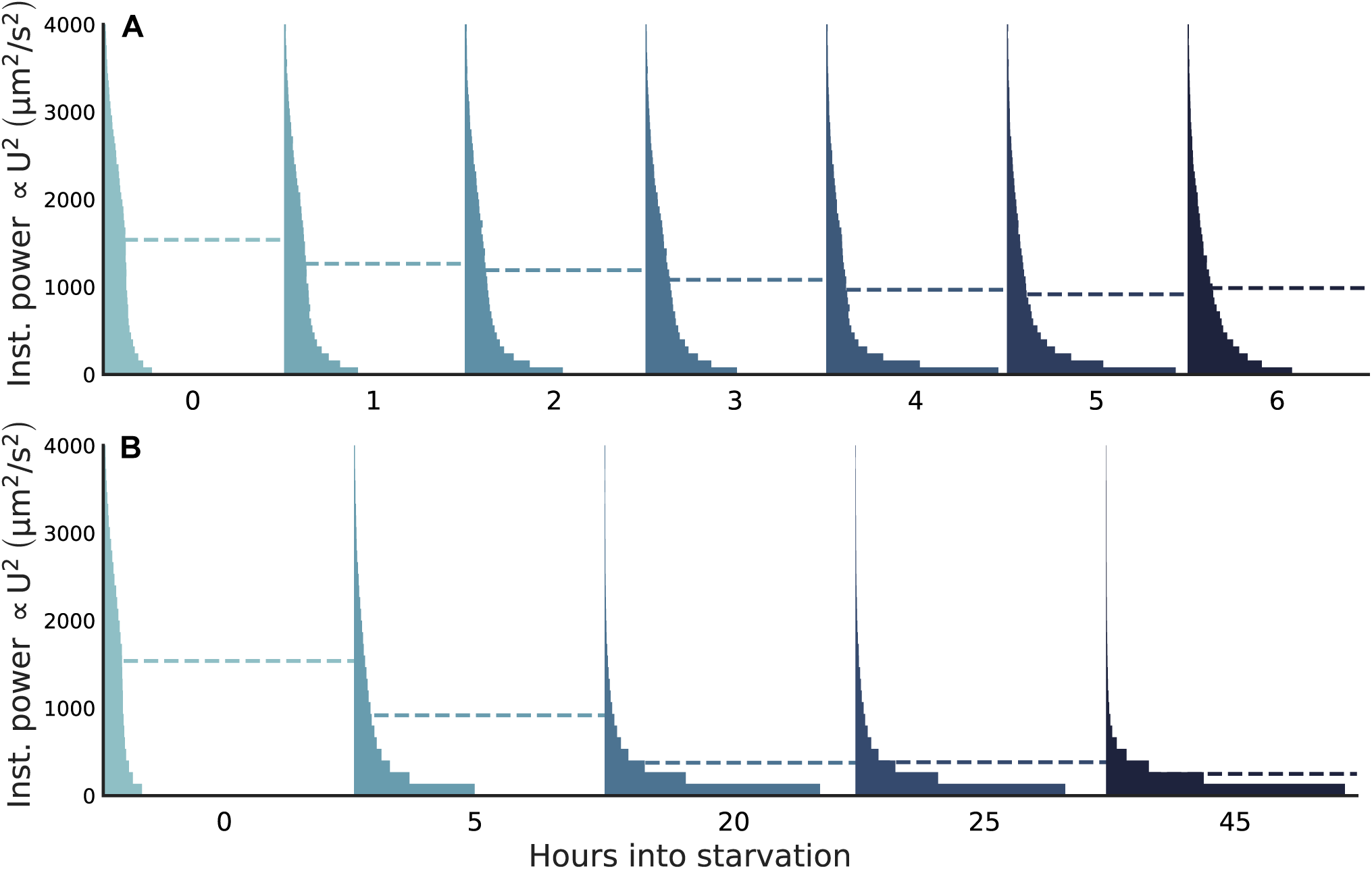
Under starvation conditions, *B. bacteriovorus* dissipates less power as it ages. From Eq Eq. (1), the power is roughly proportional to the squared speed (U^2^). (A) From Hour 0 to Hour 6, the power distribution shows a small decreasing trend. (B) From Hour 0 to Hour 45 hour, this trend becomes more apparent. The average power for each hour is noted as a dashed line. The sampling statistics for each histogram can be found in S3 Text.

Under starvation conditions, *B. bacteriovorus* expends a slightly decreasing average power (dashed line) over the first six hours. As *B. bacteriovorus* ages, the average dissipated power continues decreasing until leveling off at a constant value at which point *B. bacteriovorus* exhibits primarily diffusive behavior and the power dissipation relaxes to that predicted by the fluctuation-dissipation theorem (67). Additional runs can be found in Fig. S7, Fig. S8, and Fig. S9.

### Spiking of *B. bacteriovorus*

Following up on the VACF analysis in the previous section, we exposed previously starving bacteria to nutrient-rich media [lysogeny broth, LB], as described in the Methods. One of the main ingredients, yeast extract, has been shown to induce an active chemotactic response in *B. bacteriovorus* (11). In Fig. 5, we compare the motile response to the addition of LB to a control where *B. bacteriovorus* is kept in nutrient-poor media [HEPES media, HM], identical to those in Figs. 1 to 4.

**Fig 5.**
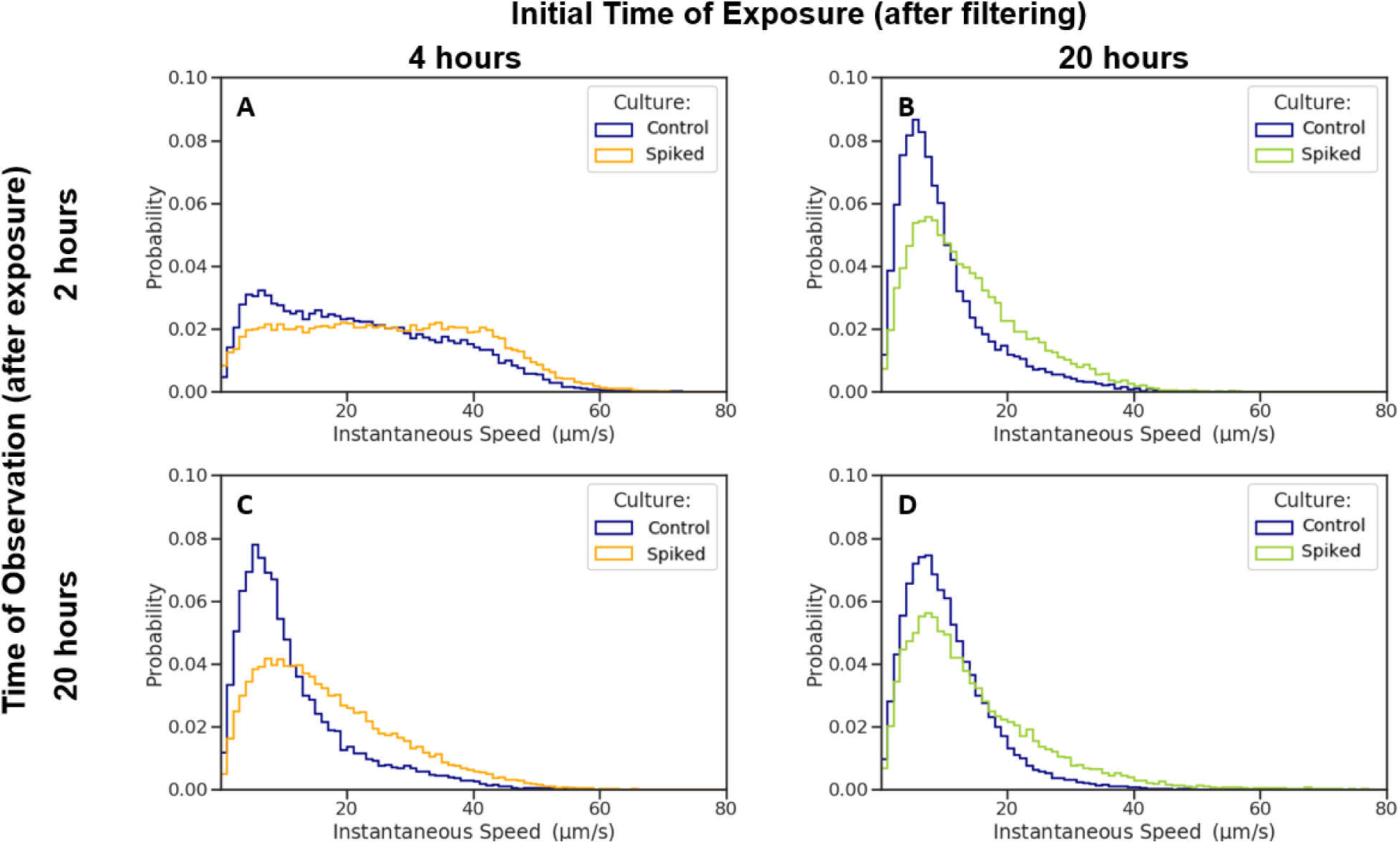
After addition of LB, *B. bacteriovorus* begins to swim faster even after long starvation times. We compare the instantaneous speed distributions of our control, starving *B. bacteriovorus* (blue), to that of cultures spiked with LB at Hour 4 (orange) and Hour 20 (green). Both spiked cultures were observed after 2 hours and 20 hours of exposure. (A) After two hours of exposure, the culture spiked at Hour 4 (orange) has a small re-population of the fast speed peak. (B) After two hours of exposure, the culture spiked at Hour 20 (green) has faster speeds than the control (blue). (C) After 20 hours of exposure, the culture spiked at Hour 4 still remains swimming faster than the control (blue). (D) After 20 hours of exposure, the culture spiked at Hour 20 still has faster instantaneous speeds than the control.The sampling statistics for each histogram can be found in S3 Text.

To determine if *B. bacteriovorus* is revivable, we took a *B. bacteriovorus* culture and separated it into three. One of these cultures was spiked with LB at Hour 4, another was spiked with LB at Hour 20, and the third was our control which was never spiked. The spiked cultures remained in the LB solution for the duration of the remainder of the experiment. For both spiked cultures, we then looked at their instantaneous speed distributions after two hours of exposure (Fig. 5A, B). We found that bacteria move slightly faster as compared to the control at the same time. On the other hand, after 20 hours of exposure (C, D), both spiked cultures at Hour 4 and 20 have higher speeds as compared to the control. Additional runs are provided in Fig. S10 and Fig. S11.

To determine if the concentration of LB used to spike the cultures at Hour 4 and 20 had an overall effect on the speed distributions, we spiked cultures at half the concentration of LB that was used to spike the cultures as well at 10X the concentration (see Fig. S12). The half LB spiked cultures exhibited similar results to the initial whole concentration seen in Fig. 5. On the other hand, at 10X LB—on account of the greater osmolarity of the solution—*B. bacteriovorus* exhibited slower speeds consistent with reports in the literature (68).

In Fig. 6, after two hours of exposure, the VACF of the culture spiked at Hour 4 more closely overlaps with its control (culture that was never spiked) in comparison to that spiked at Hour 20. However, after 20 hours of exposure to LB, the VACFs for both cultures decay *more slowly* than the control, indicating that exposure to LB induces a greater degree of *directed* motility. Taken together, these observations suggest that *B. bacteriovorus* can be revived under more favorable external conditions. Additional runs can be seen in Fig. S13 and Fig. S14.

**Fig 6.**
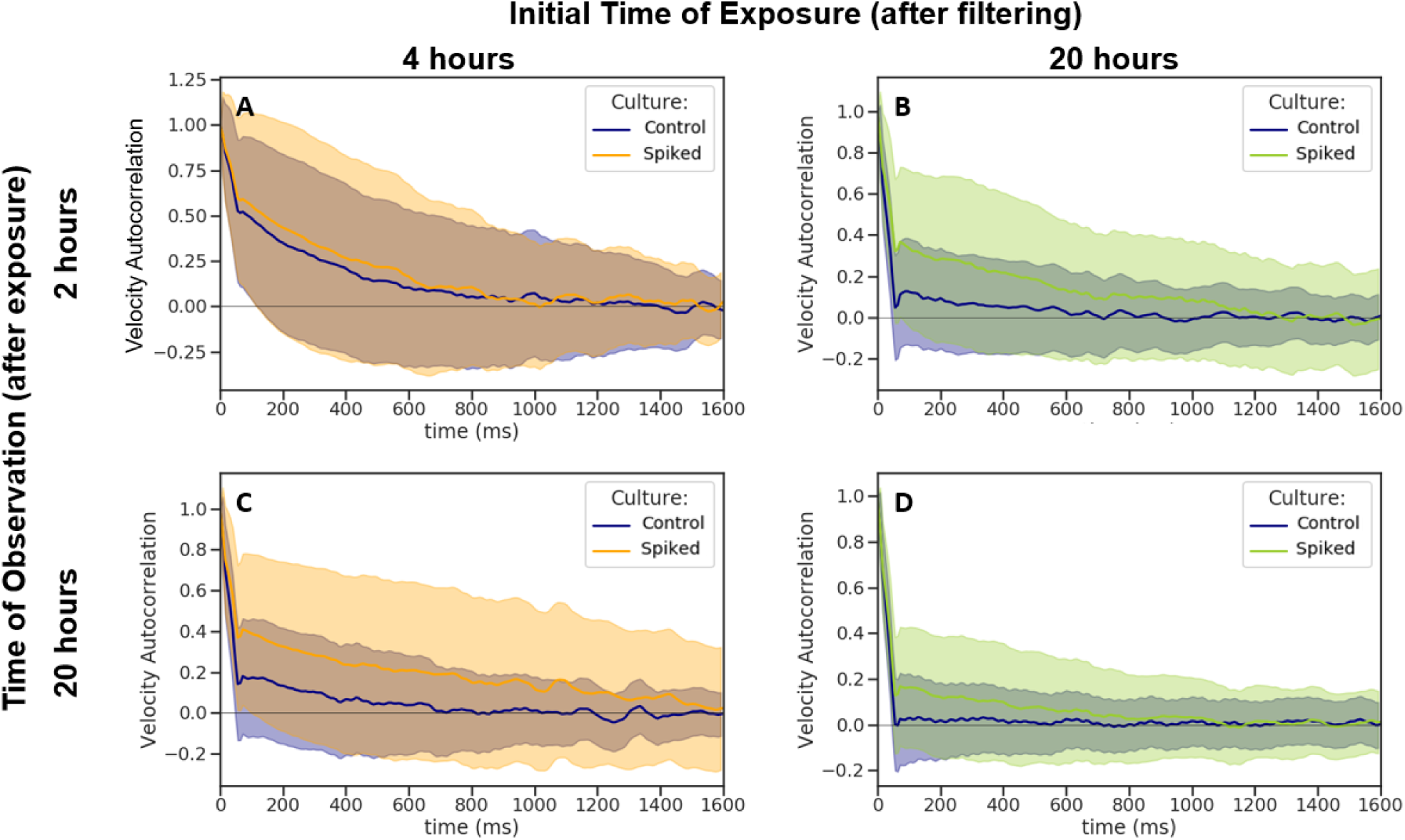
*B. bacteriovorus* shows a more slowly decaying VACF after the addition of rich media, LB. Here we compare the instantaneous speed distributions of our control, starving *B. bacteriovorus* (blue), to that of cultures spiked with LB at Hour 4 (orange) and Hour 20 (green). Both spiked cultures were observed after 2 hours and 20 hours of exposure. (A) After two hours of exposure, the culture spiked at Hour 4 (orange) decays slightly slower than the control (blue). (B) After two hours of exposure, the culture spiked at Hour 20 (green) decays slower than the control. (C) After 20 hours of exposure, the culture spiked at Hour 4 continues to decay more slowly than the control. (D) After 20 hours of exposure, the culture spiked at Hour 20 continues to decay more slowly than the control. The sampling statistics for each histogram can be found in S3 Text.

Spiking *B. bacteriovorus* at Hour 40 resulted in no change as compared to the starving control (Fig. S15 and Fig. S16). This indicates that although the speed distribution and VACF seen in Fig. 1 and Fig. 3, look similar for Hours 20 and 45, the bacteria cannot be revived after Hour 45. Therefore, it appears that *B. bacteriovorus* may strategically cease movement so as to conserve energy prior to becoming inviable.

In Fig. 7, we show the average power dissipated by starving *B. bacteriovorus* as compared to those cultures that were spiked at Hours 4 and 20. As we have shown earlier in Fig. 4, starving bacteria expend less energy as a function of their age. However, once spiked, the cultures spiked at Hour 4 (orange) and Hour 20 (green) exhibit a relatively constant amount of energy dissipated over tens of hours (e.g., from Hour 20 to Hour 45 from the start of the experiment). This suggests that, in the presence of excess nutrients, *B. bacteriovorus* expend energy at a constant rate.

**Fig 7.**
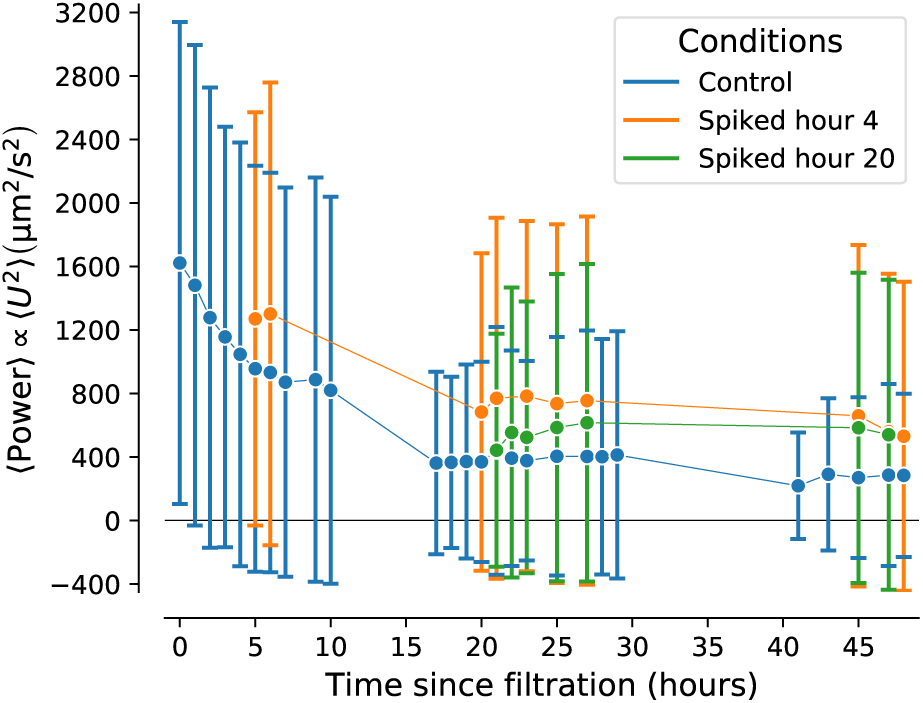
*B. bacteriovorus* dissipates more power after addition of rich media. Here we show the average dissipated power computed over all trajectories from Hour 0 to Hour 48 across multiple different runs. Our control, starving *B. bacteriovorus* (blue), is compared to cultures spiked with LB at Hour 4 (orange) and Hour 20 (green). Both spiked cultures have a larger average power after exposure to LB in comparison to our control. Over tens of hours of exposure to LB, the bacteria expend a relatively constant amount of energy. The sampling statistics for each histogram can be found in S3 Text.

Our next question was, does *B. bacteriovorus* recover its bimodal behavior following re-initialization of starvation conditions (resuspension in HM) after spiking? Here, we follow up on the spiking experiments (Fig. 5), except that after spiking we centrifuged *B. bacteriovorus* and resuspended them in nutrient-poor media. We then ask how the instantaneous speed distributions of the re-starved culture (Fig. 8B) differ from our control, starving *B. bacteriovorus* that had never been spiked (A), and that which had remained spiked (C). Qualitatively, we find that the bacteria continue to move faster as compared to the control—regardless of the tendency of centrifugation to slow down *B. bacteriovorus* (see Fig. S18)—suggesting that there is some degree of memorylessness. In other words, they behave as if they were exposed to rich media and starved for the very first time. By contrast, the re-starved culture does not swim as fast as if it had remained spiked. For further details regarding the re-initiation of starvation conditions, see the section on “Memory of *B. bacteriovorus*” in Methods.

**Fig 8.**
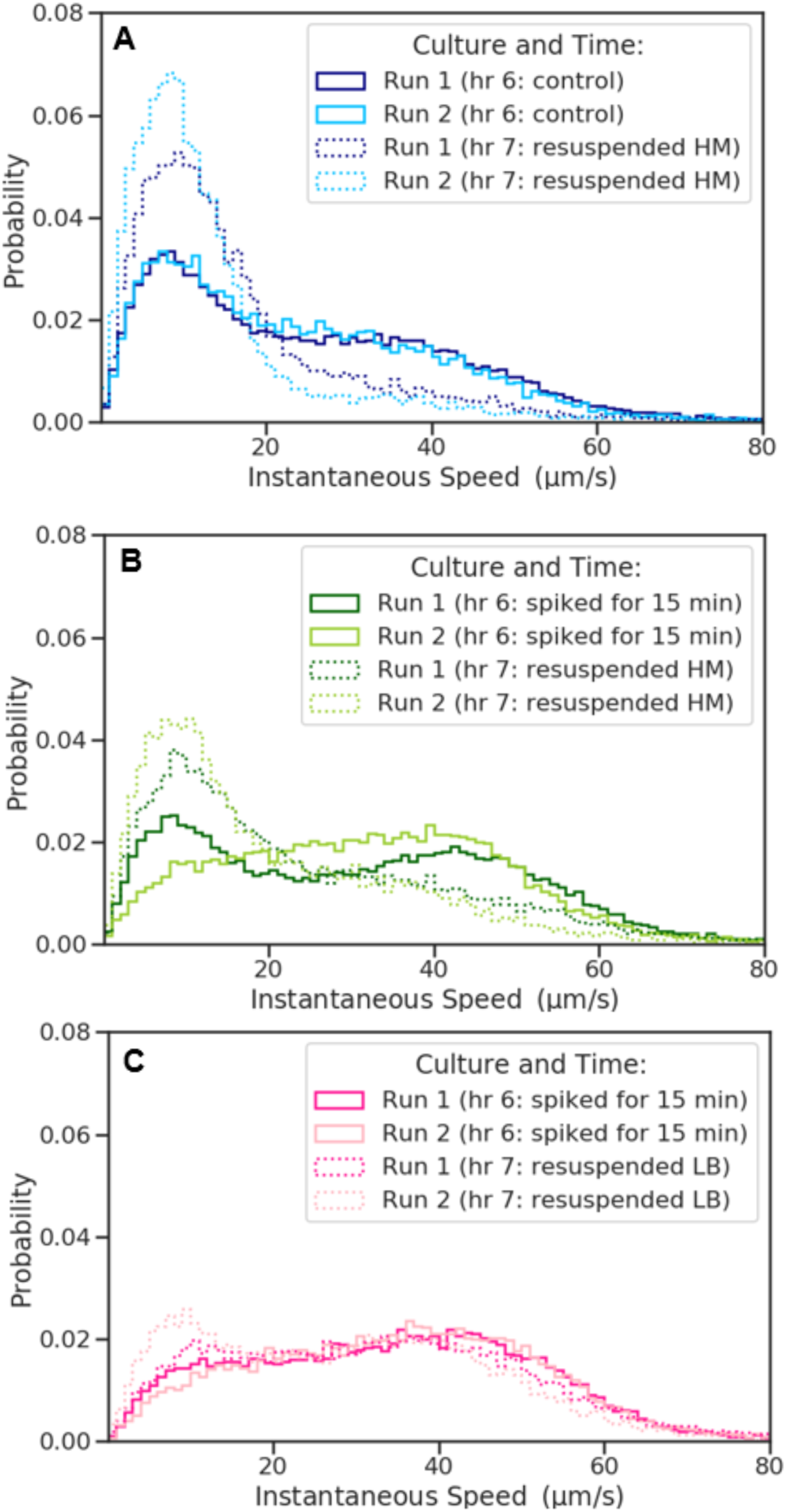
*B. bacteriovorus* decreases its speeds under starvation conditions after spiking with LB. Two runs were conducted in which each culture was starved for 6 hours then split into thirds. One of the cultures was our control, starving *B. bacteriovorus*, and two of the cultures were spiked with LB and then centrifuged and resuspended in HM (green) or spiked with LB (pink). (A) Our control culture had an increase in the height of the slow peak from Hour 6 to Hour 7 as expected. (B) The culture spiked at Hour 6 resuspended in HM then has an increase in the slow peak, but not as great of a change as seen with the control. (C) The culture spiked at Hour 6 then resuspended and spiked with LB had little to no change in the speed distribution. This indicates that *B. bacteriovorus* still decreases its speed after initiating starvation conditions (green) after being spiked, but remains swimming faster than would have been just starving. The sampling statistics for each histogram can be found in S3 Text.

### Starvation of *E. coli*

A common prey of *B. bacteriovorus* is *E. coli* (33). To compare the strategy of predatory and prey bacteria, we measured the changes in motility of *E. coli* as a function of starvation. Naively, we would expect the predators to keep moving for longer as they are always on the search for prey. In Fig. 9, we compare the instantaneous and average speed distributions of *E. coli* grown in LB (Hour -1) and after initiation of starvation conditions (Hour 0, dotted green) or resuspension in LB (Hour 0, dotted orange). The culture resuspended in LB (orange) was used as a point of comparison to quantify any bias due to centrifugation. The instantaneous and average speed distributions of this culture did not greatly vary from that taken before centrifugation.

**Fig 9.**
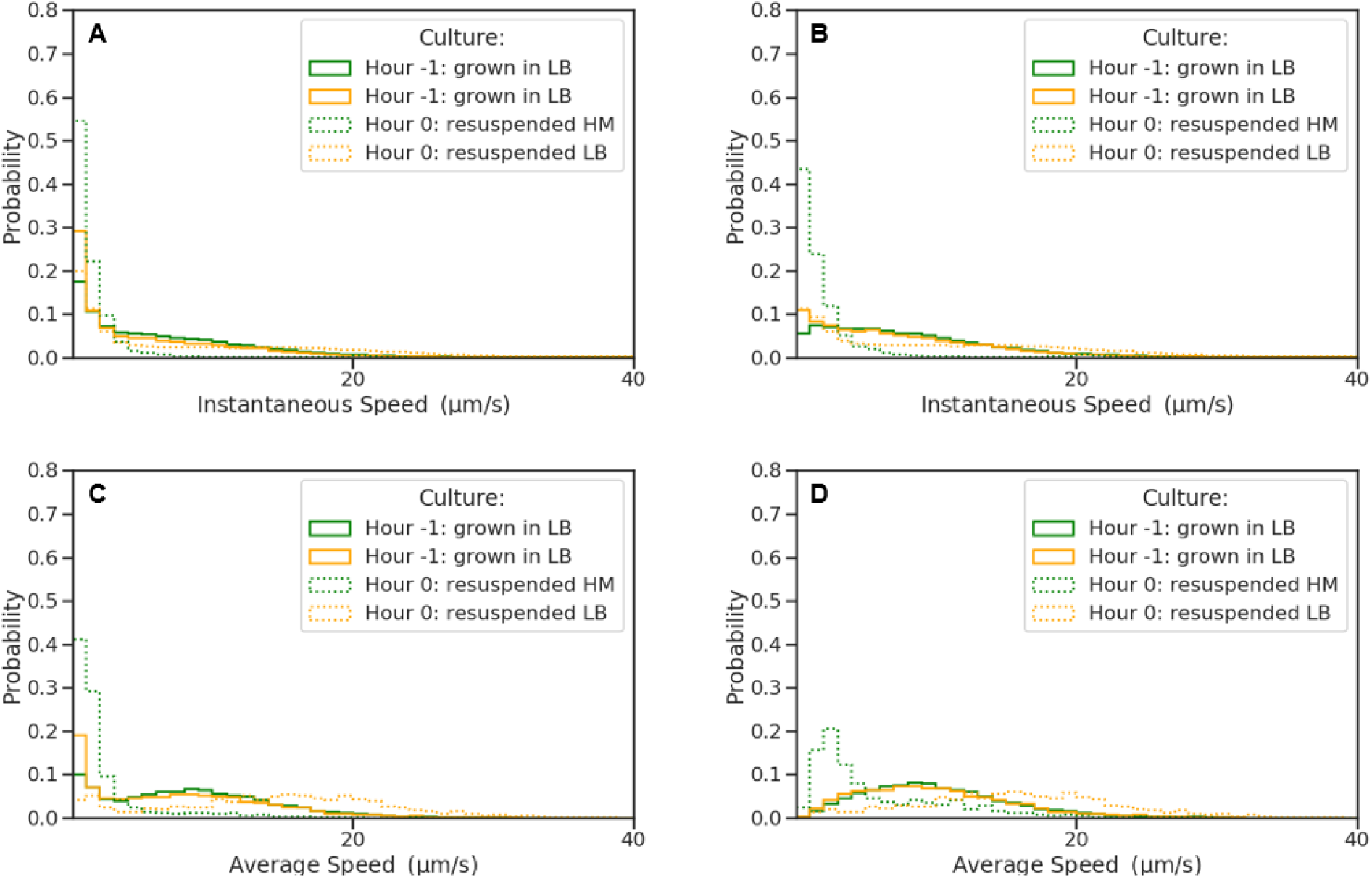
*E. coli* immediately decreases its speeds under starvation conditions. Two runs were conducted in which each *E. coli* culture was grown in LB (Hour -1) and either resuspended in HM (green) or LB (orange) at Hour 0. (A) The culture resuspended in HM (Hour 0, dotted green) had an increase in population of instantaneous slow speeds as compared to before starvation (Hour -1, solid green). (B) The motile trajectories (at least one instantaneous speed within the trajectory above 5 µm s^−1^) of *E. coli* have broader instantaneous speed distributions before centrifugation (Hour -1, solid orange and green) and after resuspension in LB (Hour 0, orange dotted) as compared to the starved culture (Hour 0, dotted green). (C) The culture resuspended in HM (Hour 0, dotted green) had an increase in population of average slow speeds as compared to prior to starvation (Hour -1, solid green) and the culture resuspended in LB (dotted orange). (D) The motile trajectories of *E. coli* have broader average speed distributions before centrifugation (Hour -1, solid orange and green) and after resuspension in LB (Hour 0, orange dotted) as compared to the starved culture (Hour 0, dotted green). The sampling statistics for each histogram can be found in S3 Text.

By contrast, after initiating starvation conditions, there are almost no motile *E. coli*, as seen with a large slow-speed population less than 5 µm s^−1^ in both the instantaneous (A) and average (C) speed distributions. The speed distributions of *E. coli* that had at least one instantaneous speed within their trajectory above 5 µm s^−1^ (i.e., motile bacteria) can be seen exclusively in Fig. 9B,D. This further indicates that even the motile fraction of starved *E. coli* has much slower speeds than before starvation.

## CONCLUSION

Bimodality in the instantaneous speed distributions over the first six hours under starvation (1A, B), along with the absence of bimodality in the trajectory-averaged speed distributions (1C, D), suggests that *B. bacteriovorus* samples both slow and fast modes of motility *within* the course of a single trajectory. At any given time-point (after the initiation of starvation), the motile behavior of a given *B. bacteriovorus* appears to be representative of other cells within the population. Indeed, *B. bacteriovorus* cells sample speeds from a bimodal distribution, with slow and fast modes, from the onset of starvation. As such, slow instantaneous speeds are not exclusively the outcome of passive diffusion. This is confirmed by representative bacterial trajectories shown in Fig. 2.

By contrast, *B. bacteriovorus’* prey, *E. coli* (Fig. 9), stop swimming under starvation conditions almost immediately, consistent with the estimated amount of time it would take for its ATP reserves to be completely depleted by constantly running its flagellar motors in nutrient poor media. In particular, the depletion time of ATP reserves from the rotating flagellum alone (excluding costs of homeostatic upkeep) can be simply estimated by considering the electrochemical potential (per proton) across the cell membrane (which is roughly 2 × 10^−20^ J (69, 70)). Considering that about 1000 protons are required per full rotation of the flagellar motor (71) under the further assumption of a modest rotational speed of 100 Hz (72, 73), we can estimate the required power as (1000 H^+^ /turn /flagellum) × (100 Hz) × (4 flagella) × (2 × 10^−20^ J) ≈10^−14^ W. Taking 10*k*_*B*_*T* for the free energy of ATP hydrolysis (74), the ATP consumption rate is roughly 2.5 ×10^6^ ATP s. Taking the volume of an *E. coli* cell to be ≈1 µm^3^, the ATP concentration to be 1.54 mm (75), and the ATP consumption rate of the flagellar motor to be 2.5 × 10^6^ ATP s, we find that the flagellar motor of *E. coli* can in principle consume all available ATP in the cell in well under a second in the absence of new ATP production.

By contrast, remarkably, *B. bacteriovorus* remains active over hours and our spiking experiments not only show that *B. bacteriovorus* can be resuscitated after even 20 hours of starvation, but that those bacteria that *do* stop swimming (and revert to passive diffusion) can be stimulated back into sampling the high speed mode of its velocity distribution. Indeed, shortly after spiking, *B. bacteriovorus* substantially increases its swim speed and shows a greater directed motion (as measured by the VACF, Fig. 3). As the re-initiation of starvation conditions (Fig. 8), produced behavior similar to initial starvation, it is likely that *B. bacteriovorus* allocates energy reserves for active motion in direct response to its environment, regardless of previous conditions, and that the *B. bacteriovorus* population eventually enters an energy-conserving dormant stage if starved long enough. Previous studies in which bimodal speed distributions are observed are normally associated to the bundling of flagella for peritrichously flagellated species (e.g., the run-and-tumble paradigm (76, 77)) which is not a feasible mechanism for the uniflagellated *B. bacteriovorus*.

The subtle way in which *B. bacteriovorus* modulates which of the modes of its speed distribution it samples is worthy, in and of itself, of future molecular attention. To wit, it is already known that *B. bacteriovorus* dramatically varies its gene expression levels between attack and growth phases (59, 78, 79) with, most recently, work elucidating *B. bacteriovorus*’s motility reduction under starvation linked to cyclic-di-GMP effectors playing the role of “motility brakes” (80).

These findings, in turn, beg broader questions associated to the cost, both hydrodynamic and transcriptional, tied to motility re-initiation upon detection of environmental cues by *B. bacteriovorus*. Our work here provides a piece of the puzzle toward assessing the cost associated to motility with age. It remains to be seen whether, in the end, complex speed modulation with age is a by-product of varying transcriptional activity or part of a global strategy to jointly minimize hydrodynamic and transcriptional energy expenditure in order to maximize the probability of locating prey.

## AUTHOR CONTRIBUTIONS

Conceived and designed the experiments: SP MC. Performed the experiments: MC. Analyzed the data: MC SLS. Contributed reagents/materials/analysis tools: SLS. Wrote and edited the paper: MC SLS SP. Designed the software: SLS MC. Oversaw every aspect of the research: SP.

## ACKNOWLEDGMENTS

S. Pressé acknowledges the National Institute of Health (NIH under grant No. AWD00033604) and the National Science Foundation Career (NSF under grant No. GR02240). We would like to thank Doug Shepherd for helpful feedback on all experimental aspects of this study. We would also like to give a special thanks to Edouard Jurkevitch for his help with *B. bacteriovorus* growth protocols and Yixin Shi for generously providing *E. coli* strains.

## Notes

### Competing Interest Statement

The authors have declared no competing interest.

## REFERENCES

1. Drescher, K., J. Dunkel, L. Cisneros, S. Ganguly, and R. Goldstein, 2011. Fluid dynamics and noise in bacterial cell-cell and cell-surface scattering. Proc. Nat. Acad. Sci. 108:10940.

2. Jashnsaz, H., M. Al Juboori, C. Weistuch, N. Miller, T. Nguyen, V. Meyerhoff, B. McCoy, S. Perkins, R. Wallgren, B. D. Ray, K. Tsekouras, G. G. Anderson, and S. Pressé, 2017. Hydrodynamic Hunters. Biophys. J. 112:1282–1289.

3. Meng, Y., Y. Li, C. Galvani, G. Hao, J. Turner, T. Burr, and H. Hoch, 2005. Upstream migration of *Xylella fastidiosa* via pilus-driven twitching motility. J. Bacteriol. 187:5560.

4. Shen, Y., A. Siryaporn, S. Lecuyer, Z. Gitai, and H. Stone, 2012. Flow directs surface-attached bacteria to twitch upstream. Biophys. J. 103:146.

5. Kaya, T., and H. Koser, 2012. Direct upstream motility in *Escherichia coli*. Biophys. J. 102:1514.

6. Baker, M., P. Wolanin, and J. Stock, 2006. Signal transduction in bacterial chemotaxis. BioEssays 28:9.

7. Dori-Bachash, M., B. Dassa, S. Pietrokovski, and E. Jurkevitch, 2008. Proteome-based comparative analyses of growth stages reveal new cell cycle-dependent functions in the predatory bacterium *Bdellovibrio bacteriovorus*. Appl. Environ. Microbiol. 74:7152.

8. Waters, C., and B. Bassler, 2005. Quorum Sensing: Cell-to-Cell Communication in Bacteria. Annu. Rev. Cell Dev. Biol. 21:319.

9. Mukherjee, S., S.-C. Seok, V. Vieland, and J. Das, 2013. Data-driven quantification of the robustness and sensitivity of cell signaling networks. Phys. Biol. 10:066002.

10. Jiang, L., Q. Ouyang, and Y. Tu, 2010. Quantitative Modeling of *Escherichia coli* Chemotactic Motion in Environments Varying in Space and Time. PLoS Comput. Biol. 6:e1000735.

11. LaMarre, A., S. Straley, and S. Conti, 1977. Chemotaxis toward amino acids by *Bdellovibrio bacteriovorus*. Journal of bacteriology 131:201–207.

12. Hespell, R. B., M. F. Thomashow, and S. C. Rittenberg, 1974. Changes in cell composition and viability of *Bdellovibrio bacteriovorus* during starvation. Arch. Microbiol. 97:313–327.

13. Dashiff, A., R. Junka, M. Libera, and D. Kadouri, 2011. Predation of human pathogens by the predatory bacteria *Micavibrio aeruginosavorus* and *Bdellovibrio bacteriovorus*. J. Appl. Microbiol. 110:431.

14. Sockett, R., and C. Lambert, 2004. *Bdellovibrio* as therapeutic agents: a predatory renaissance? Nat. Rev. Micro. 2:669.

15. Kadouri, D., and G. O’Toole, 2005. Susceptibility of biofilms to *Bdellovibrio bacteriovorus* attack. Appl. Environ. Microbiol. 71:4044.

16. Passino, K., 2002. Biomimicry of bacterial foraging for distributed optimization and control. Control Syst. Mag. 22:52.

17. Astling, D., J. Lee, and D. Zusman, 2006. Differential effects of chemoreceptor methylation-domain mutations on swarming and development in the social bacterium *Myxococcus xanthus*. Mol. Microbiol. 59:45.

18. Williams, H., J. Kelley, M. Baer, and B. Turng, 1995. The association of *Bdellovibrios* with surfaces in the aquatic environment. Can. J. Microbiol. 41:1142.

19. Lambert, C., A. Fenton, L. Hobley, and R. Sockett, 2011. Predatory *Bdellovibrio* Bacteria Use Gliding Motility To Scout for Prey on Surfaces. J. Bacteriol. 193:139.

20. Harini, K., V. Ajila, and S. Hegde, 2013. Bdellovibrio bacteriovorus: A future antimicrobial agent?

21. Kadouri, D., and G. A. O’Toole, 2005. Susceptibility of biofilms to *Bdellovibrio bacteriovorus* attack. Appl. Environ. Microbiol. 71:4044–4051.

22. Chanyi, R. M., and S. F. Koval, 2014. Role of Type IV Pili in Predation by *Bdellovibrio bacteriovorus*. PLoS One 9.

23. Straley, S., and S. Conti, 1977. Chemotaxis by *Bdellovibrio bacteriovorus* toward prey. J. Bacteriol. 132:628.

24. Oyedara, O., D. Luna-Santillana, E. de Jesus, O. Olguin-Rodriguez, X. Guo, M. Mendoza-Villa, J. Menchaca-Arredondo, T. Elufisan, J. Garza-Hernandez, I. Garcia Leon, et al., 2016. Isolation of *Bdellovibrio sp*. from soil samples in Mexico and their potential applications in control of pathogens. MicrobiologyOpen 5:992.

25. Yu, R., S. Zhang, Z. Chen, and C. Li, 2017. Isolation and application of predatory *Bdellovibrio*-and-like organisms for municipal waste sludge biolysis and dewaterability enhancement. Front. Env. Sci. Eng. 11:10.

26. Sun, Y., J. Ye, Y. Hou, H. Chen, J. Cao, and T. Zhou, 2017. Predation Efficacy of *Bdellovibrio bacteriovorus* on Multidrug-Resistant Clinical Pathogens and Their Corresponding Biofilms. Jpn. J. Infect. Dis. 70:485–489.

27. Iebba, V., V. Totino, F. Santangelo, A. Gagliardi, L. Ciotoli, A. Virga, C. Ambrosi, M. Pompili, R. V. De Biase, L. Selan, M. Artini, F. Pantanella, F. Mura, C. Passariello, M. Nicoletti, L. Nencioni, M. Trancassini, S. Quattrucci, and S. Schippa, 2014. Bdellovibrio bacteriovorus directly attacks Pseudomonas aeruginosa and Staphylococcus aureus cystic fibrosis isolates. Front. Microbiol. 5.

28. Shatzkes, K., E. Singleton, C. Tang, M. Zuena, S. Shukla, S. Gupta, S. Dharani, O. Onyile, J. Rinaggio, N. Connell, and D. Kadouri, 2016. Predatory Bacteria Attenuate *Klebsiella pneumoniae* Burden in Rat Lungs. MBio. 7:e01847.

29. Atterbury, R., L. Hobley, R. Till, C. Lambert, M. Capeness, T. Lerner, A. Fenton, P. Barrow, and R. Sockett, 2011. Effects of orally administered *Bdellovibrio bacteriovorus* on the well-being and *Salmonella* colonization of young chicks. Appl. Environ. Microbiol. 77:5794.

30. Romanowski, E., N. Stella, K. Brothers, K. Yates, M. Funderburgh, J. Funderburgh, S. Gupta, S. Dharani, D. Kadouri, and R. Shanks, 2016. Predatory bacteria are nontoxic to the rabbit ocular surface. Sci. Rep. 6:30987.

31. Willis, A., C. Moore, M. Mazon-Moya, S. Krokowski, C. Lambert, R. Till, S. Mostowy, and R. Sockett, 2016. Injections of Predatory Bacteria Work Alongside Host Immune Cells to Treat *Shigella* Infection in Zebrafish Larvae. Curr. Biol. 26:3343.

32. Park, S., D. Kim, R. J. Mitchell, and T. Kim, 2011. A microfluidic concentrator array for quantitative predation assays of predatory microbes. Lab Chip 11:2916–2923.

33. Van Essche, M., M. Quirynen, I. Sliepen, J. Van Eldere, and W. Teughels, 2009. *Bdellovibrio bacteriovorus* attacks *Aggregatibacter actinomycetemcomitans*. J. Dent. Res. 88:182–186.

34. Mitchell, R., S. Yankfsky, and H. W. Jannasch, 1967. Lysis of *Escherichia coli* by marine micro-organisms. Nature 215:891–893.

35. Lambert, C., M. C. M. Smith, and R. E. Sockett, 2003. A novel assay to monitor predator-prey interactions for *Bdellovibrio bacteriovorus* 109 J reveals a role for methyl-accepting chemotaxis proteins in predation. Environ. Microbiol. 5:127–132.

36. Rendulic, S., P. Jagtap, A. Rosinus, M. Eppinger, C. Baar, C. Lanz, H. Keller, C. Lambert, K. J. Evans, A. Goesmann, F. Meyer, R. E. Sockett, and S. C. Schuster, 2004. A predator unmasked: life cycle of *Bdellovibrio bacteriovorus* from a genomic perspective. Science 303:689–692.

37. Steyert, S. R., and S. A. Pineiro, 2007. Development of a novel genetic system to create markerless deletion mutants of *Bdellovibrio bacteriovorus*. Appl. Environ. Microbiol. 73:4717–4724.

38. Lambert, C., K. J. Evans, R. Till, L. Hobley, M. Capeness, S. Rendulic, S. C. Schuster, S. I. Aizawa, and R. E. Sockett, 2006. Characterizing the flagellar filament and the role of motility in bacterial prey-penetration by *Bdellovibrio bacteriovorus*. Mol. Microbiol..

39. Lambert, C., K. A. Morehouse, C. Y. Chang, and R. E. Sockett, 2006. Bdellovibrio: growth and development during the predatory cycle.

40. Thomashow, M. F., and T. W. Cotter, 1992. *Bdellovibrio* host dependence: the search for signal molecules and genes that regulate the intraperiplasmic growth cycle. J. Bacteriol. 174:5767–5771.

41. Fratamico, P. M., and R. C. Whiting, 1995. Ability of *Bdellovibrio bacteriovorus* 109J to Lyse Gram-Negative Food-Borne Pathogenic and Spoilage Bacteria. J. Food Prot. 58:160–164.

42. Seidler, R. J., and M. P. Starr, 1969. Factors affecting the intracellular parasitic growth of *Bdellovibrio bacteriovorus* developing within Escherichia coli. Journal of Bacteriology 97:912–923.

43. Stolp, H., and H. Petzold, 1962. Untersuchungen ueber einen obligat parasitishen Mikroorganismus mit lyticher Aktivitaet fuer Pseudomonas-bakterien. Phytopathology 45:364.

44. Stolp, H., and M. Starr, 1963. *Bdellovibrio bacteriovorus gen. et sp. n.*, a predatory, ectoparasitic, and bacteriolytic microorganism. Antonie Van Leeuwenhoek 29:217.

45. Seidler, R., and M. Starr, 1969. Isolation and Characterization of Host-Independent *Bdellovibrios*. J. Bacteriol. 100:769.

46. Lambert, C., M. Smith, and R. Sockett, 2003. A novel assay to monitor predator-prey interactions for *Bdellovibrio bacteriovorus* 109J reveals a role for methyl-accepting chemotaxis proteins in predation. Environ. Microbiol. 5:127.

47. Varon, M., and B. Zeigler, 1978. Bacterial predator-prey interaction at low prey density. Appl. Environ. Microbiol. 36:11.

48. Lambert, C., K. Morehouse, C. Chang, and R. Sockett, 2006. *Bdellovibrio*: growth and development during the predatory cycle. Curr. Opin. Microbiol. 9:639.

49. Lambert, C., L. Hobley, C. Chang, A. Fenton, M. Capeness, and L. Sockett, 2008. A predatory patchwork: membrane and surface structures of *Bdellovibrio bacteriovorus*. Adv. Microb. Physiol. 54:313.

50. Straley, S., and S. Conti, 1974. Chemotaxis by *Bdellovibrio bacteriovorus*. J. Bacteriol. 120:549.

51. Strauch, E., D. Schwudke, and M. Linscheid, 2007. Predatory mechanisms of *Bdellovibrio* and like organisms. Future Microbiol. 2:63.

52. Drescher, K., R. Goldstein, N. Michel, M. Polin, and I. Tuval, 2010. Direct Measurement of the Flow Field around Swimming Microorganisms. Phys. Rev. Lett. 105:168101.

53. Lushi, E., H. Wioland, and R. Goldstein, 2014. Fluid flows created by swimming bacteria drive self-organization in confined suspensions. Proc. Natl. Acad. Sci. 111:9733.

54. Frymier, P., R. Ford, H. Berg, and P. Cummings, 1995. Three-dimensional tracking of motile bacteria near a solid planar surface. Proc. Nat. Acad. Sci. 92:6195.

55. Lauga, E., W. DiLuzio, G. Whitesides, and H. Stone, 2006. Swimming in Circles: Motion of Bacteria near Solid Boundaries. Biophys. J. 90:400.

56. Di Leonardo, R., D. Dell’Arciprete, L. Angelani, and V. Iebba, 2011. Swimming with an Image. Phys. Rev. Lett. 106:038101.

57. Hu, J., A. Wysocki, R. G. Winkler, and G. Gompper, 2015. Physical Sensing of Surface Properties by Microswimmers: Directing Bacterial Motion via Wall Slip. Sci. Rep. 5:9586.

58. Guo, M. S., and C. A. Gross, 2014. Stress-induced remodeling of the bacterial proteome. Curr. Biol. 24:R424–34.

59. Karunker, I., O. Rotem, M. Dori-Bachash, E. Jurkevitch, and R. Sorek, 2013. A global transcriptional switch between the attack and growth forms of Bdellovibrio bacteriovorus. PLoS One 8:e61850.

60. Dori-Bachash, M., B. Dassa, S. Pietrokovski, and E. Jurkevitch, 2008. Proteome-based comparative analyses of growth stages reveal new cell cycle-dependent functions in the predatory bacterium Bdellovibrio bacteriovorus. Appl. Environ. Microbiol. 74:7152–7162.

61. Lambert, C., T. R. Lerner, N. K. Bui, H. Somers, S.-I. Aizawa, S. Liddell, A. Clark, W. Vollmer, A. L. Lovering, and R. E. Sockett, 2016. Interrupting peptidoglycan deacetylation during *Bdellovibrio* predator-prey interaction prevents ultimate destruction of prey wall, liberating bacterial-ghosts. Sci. Rep. 6:26010.

62. Allan, D., C. van der Wel, N. Keim, T. A. Caswell, D. Wieker, R. Verweij, C. Reid, Thierry, L. Grueter, K. Ramos, apiszcz, zoeith, R. W. Perry, F. Boulogne, P. Sinha, pfigliozzi, N. Bruot, L. Uieda, J. Katins, H. Mary, and A. Ahmadia, 2019. soft-matter/trackpy: Trackpy v0.4.2.

63. Amsler, C. D., M. Cho, and P. Matsumura, 1993. Multiple factors underlying the maximum motility of *Escherichia coli* as cultures enter post-exponential growth. J. Bacteriol. 175:6238–6244.

64. Maeda, K., Y. Imae, J. I. Shioi, and F. Oosawa, 1976. Effect of temperature on motility and chemotaxis of *Escherichia coli*. J. Bacteriol. 127:1039–1046.

65. Boehm, A., M. Kaiser, H. Li, C. Spangler, C. A. Kasper, M. Ackermann, V. Kaever, V. Sourjik, V. Roth, and U. Jenal, 2010. Second messenger-mediated adjustment of bacterial swimming velocity. Cell 141:107–116.

66. Magariyama, Y., S. Sugiyama, K. Muramoto, I. Kawagishi, Y. Imae, and S. Kudo, 1995. Simultaneous measurement of bacterial flagellar rotation rate and swimming speed. Biophys. J. 69:2154–2162.

67. Kubo, R., 1966. The fluctuation-dissipation theorem. Rep. Prog. Phys. 29:255.

68. Im, H., S. Son, R. J. Mitchell, and C.-M. Ghim, 2017. Serum albumin and osmolality inhibit *Bdellovibrio bacteriovorus* predation in human serum. Sci. Rep. 7:5896.

69. Kashket, E. R., 1985. The proton motive force in bacteria: a critical assessment of methods. Annu. Rev. Microbiol. 39:219–242.

70. Gabel, C. V., and H. C. Berg, 2003. The speed of the flagellar rotary motor of Escherichia coli varies linearly with protonmotive force. Proc. Natl. Acad. Sci. U. S. A. 100:8748–8751.

71. Meister, M., G. Lowe, and H. C. Berg, 1987. The proton flux through the bacterial flagellar motor. Cell 49:643–650.

72. Berg, H. C., and L. Turner, 1993. Torque generated by the flagellar motor of Escherichia coli. Biophys. J. 65:2201–2216.

73. Berry, R. M., and H. C. Berg, 1999. Torque generated by the flagellar motor of Escherichia coli while driven backward. Biophys. J. 76:580–587.

74. Rosing, J., and E. C. Slater, 1972. The value of ΔG for the hydrolysis of ATP. Biochimica et Biophysica Acta (BBA)-Bioenergetics 267:275–290.

75. Yaginuma, H., S. Kawai, K. V. Tabata, K. Tomiyama, A. Kakizuka, T. Komatsuzaki, H. Noji, and H. Imamura, 2014. Diversity in ATP concentrations in a single bacterial cell population revealed by quantitative single-cell imaging. Scientific reports 4:6522.

76. Toley, B. J., and N. S. Forbes, 2012. Motility is critical for effective distribution and accumulation of bacteria in tumor tissue. Integrative Biology 4:165–176.

77. Matz, C., and K. Jürgens, 2005. High motility reduces grazing mortality of planktonic bacteria. Applied and Environmental Microbiology 71:921–929.

78. Waso, M., S. Khan, W. Ahmed, and W. Khan, 2020. Expression of attack and growth phase genes of Bdellovibrio bacteriovorus in the presence of Gram-negative and Gram-positive prey. Microbiol. Res. 235:126437.

79. Nogales, J., C. Herencias, and M. A. Prieto, 2020. Providing new insights on the byphasic lifestyle of the predatory bacterium *Bdellovibrio bacteriovorus* through genome-scale metabolic modeling. bioRxiv.

80. Sathyamoorthy, R., Y. Kushmaro, O. Rotem, O. Matan, D. E. Kadouri, A. Huppert, and E. Jurkevitch, 2020. To hunt or to rest: prey depletion induces a novel starvation survival strategy in bacterial predators. The ISME Journal 1–15.

