## Supplemental Information for "Swimming, fast and slow: strategy and survival of bacterial predators in response to chemical cues"

### SUPPLEMENTARY MATERIAL

**S1 Text. Diffusive speed calculations.** Estimating *B. bacteriovorus* as a sphere, we can calculate the diffusion coefficient given the Stokes-Einstein equation

$$D = kT/6\pi \nu r, \quad (2)$$

in which $\nu$ is the dynamic viscosity and $r$ the radius of the sphere. The dynamic viscosity of water at 28°C (temperature of the incubator) is approximately $8.318 \times 10^{-4}$ kg m s$^{-1}$. The radius of the bacterium is approximated as 0.375 µm. By substituting these values and Boltzmann's constant, $k$, the diffusion coefficient can be estimated as 0.707 µm$^2$ s$^{-1}$. The exposure time $\Delta t$vv between two frames is 0.008 s.

$$v_{\text{diff}} = \sqrt{D/\Delta t}. \quad (3)$$

Therefore, the diffusive speed is approximately 9.40 µm s$^{-1}$. A rough calculation for *E. coli*'s diffusive speed can be obtained by approximating the cell as sphere and taking its volume to be ≈ 1 µm$^3$, giving a radius of 0.6 µm. From Eq. (3) above, we get $v_{\text{diff}} \approx 7.4$ µm s$^{-1}$.

**S2 Text. VACF calculations.** Using the instantaneous velocities for each bacterium's trajectory, the VACF was calculated as

$$Z = \frac{\langle \mathbf{v}(t) \cdot \mathbf{v}(t+\tau) \rangle}{\langle \mathbf{v}(t) \cdot \mathbf{v}(t) \rangle}, \quad (4)$$

where $\mathbf{v}(t)$ is the velocity at some time t and $\mathbf{v}(t+\tau)$ is some time later (8 ms between frames). To simplify, $\langle \mathbf{v}(t) \cdot \mathbf{v}(t) \rangle$ is equivalent to $\langle v_x(t)v_x(t) + v_y(t)v_y(t) \rangle$ given the x-y components in a particular frame of a bacterium's trajectory.

**S3 Text. Sampling statistics for main text figures.** Fig 1: The area of each histogram is normalized to unity. For (A) and (B), the number of instantaneous speed observations ranges from 65,795-151,604; for (C) and (D), the number of average speed observations ranges from 621-2,589. Instantaneous speed distributions were constructed from 50 evenly spaced bins over the range of instantaneous speed measurements; average speed distributions were constructed from 30 evenly spaced bins over the range of average speed measurements.

Fig 3: Details of VACF calculations are provided in Eq. (4).

Fig 4: The area of each histogram is normalized to unity. For (A) and (B), the number of U$^2$ observations ranges from 65,795-151,604. Distributions were constructed from 50 evenly spaced bins over the range of measured power dissipation.

Fig 5: The area of each histogram is normalized to unity. The number of instantaneous speed observations ranges from 39,244-39,862 for (A), 29,943-30,707 for (B), 18,841-20,320 for (C), and 56,115-60,393 for (D). Distributions were constructed from 100 evenly spaced bins over the range of instantaneous speed measurements.

Fig 7: The error shown for each hour is one standard deviation above and below the mean. For the control culture, Hour 0 through Hour 6, Hour 25, and Hour 27 were obtained from four runs, results for Hour 47 from three runs, and all other hours from a single run. For the culture spiked at Hour 4, Hour 6, 25, and 27 were obtained from three runs, Hours 5 and 47 from two runs, and all additional times were from one run. The number of runs contributing to the culture spiked at hour 20 was two for hours 25, 27 and 47, and one for all extra hours.

Fig 8 The area of each histogram is normalized to unity. The number of instantaneous speed observations ranges from 11,488-30,349 for (A), 12,092-43,594 for (B), and 14,292-33,060 for (C). Distributions were constructed from 50 evenly spaced bins over the range of instantaneous speed measurements.

Fig 9: The area of each histogram is normalized to unity. The number of instantaneous speed observations ranges from 117,565-345,396 for (A) and 100,263-276,050; The number of average speed observations ranges from 1,407-9,402 for (C) and 684-7.727 for (D). Instantaneous speed distributions were constructed from 50 evenly spaced bins over the range of instantaneous speed measurements; average speed distributions were constructed from 30 evenly spaced bins over the range of average speed measurements.

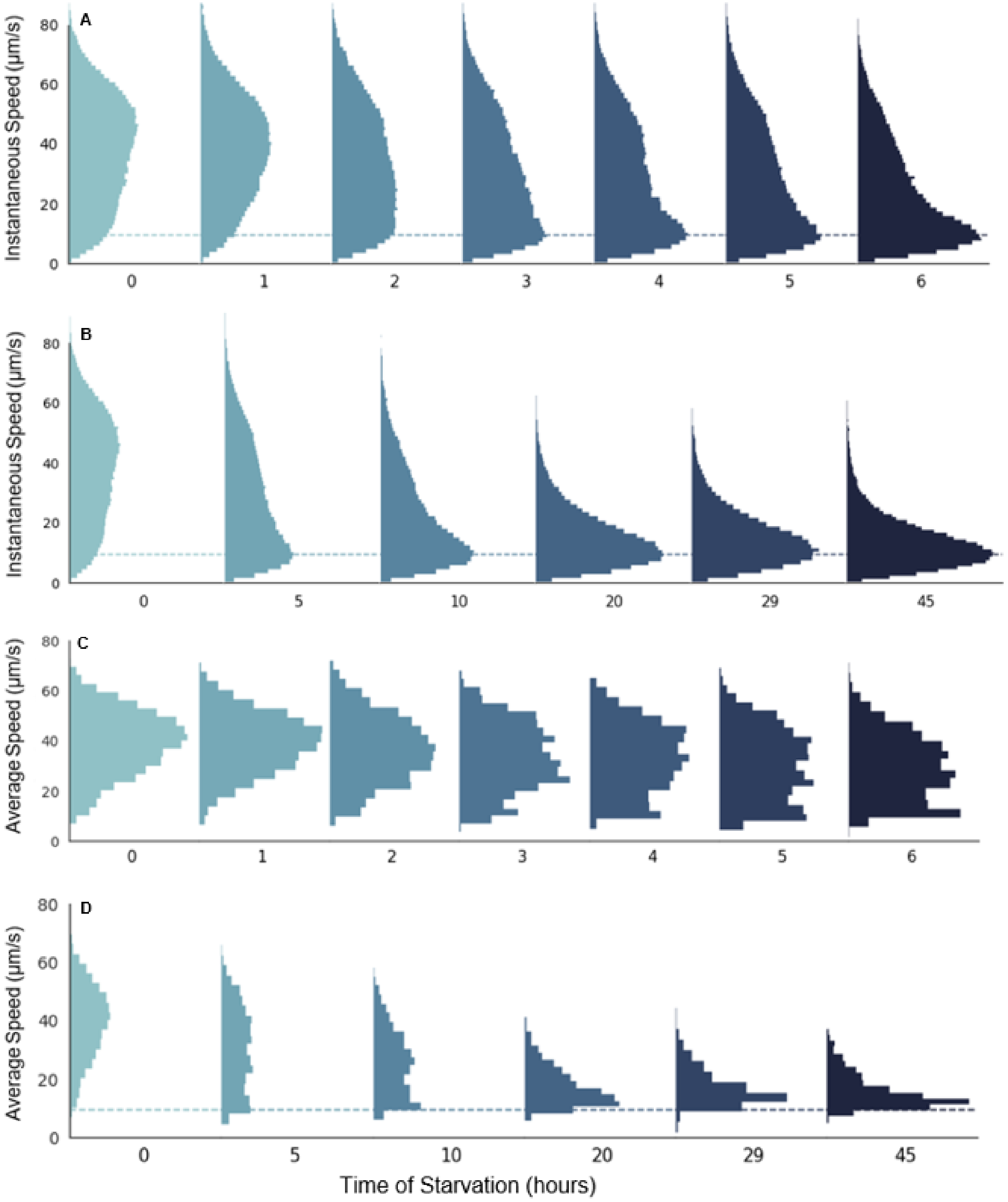


**Fig S1. Under starvation conditions, the instantaneous and average speed distributions of *B. bacteriovorus* shift across time. [run 2].** We follow a similar convention to Fig. 1 in the main text. The only relevant difference is the number of data points contributing to each histogram. For (A) and (B), the number of instantaneous speed observations ranges from 93,196-172,376; for (C) and (D), the number of average speed observations ranges from 1,327-3,277.

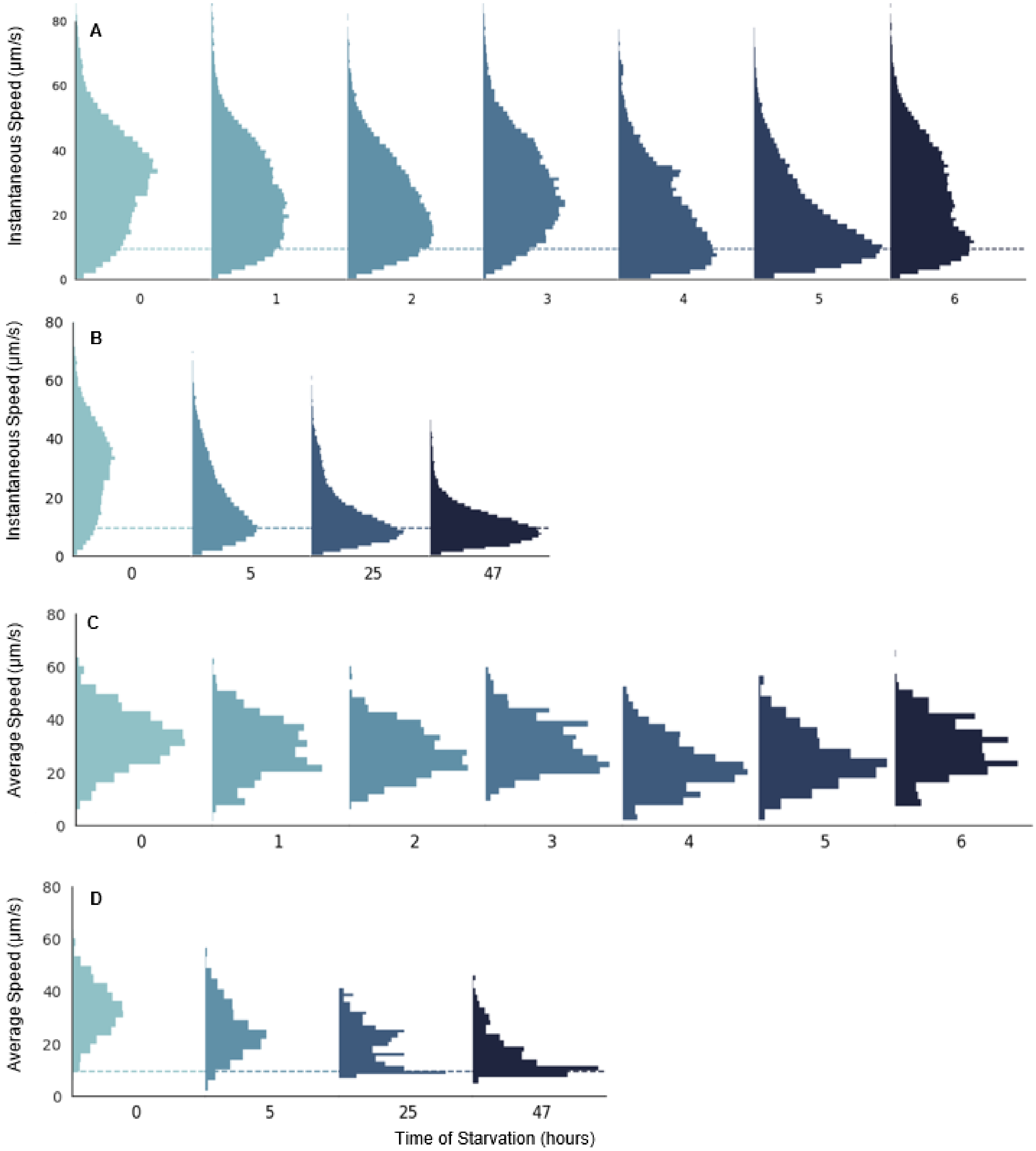


**Fig S2. Under starvation conditions, the instantaneous and average speed distributions of *B. bacteriovorus* shift across time [run 3].** We follow a similar convention to Fig. 1 in the main text. The only relevant difference is the the number of data points contributing to each histogram. For (A) and (B), the number of instantaneous speed observations ranges from 25,959-60,393; for (C) and (D), the number of average speed observations ranges from 247-982.

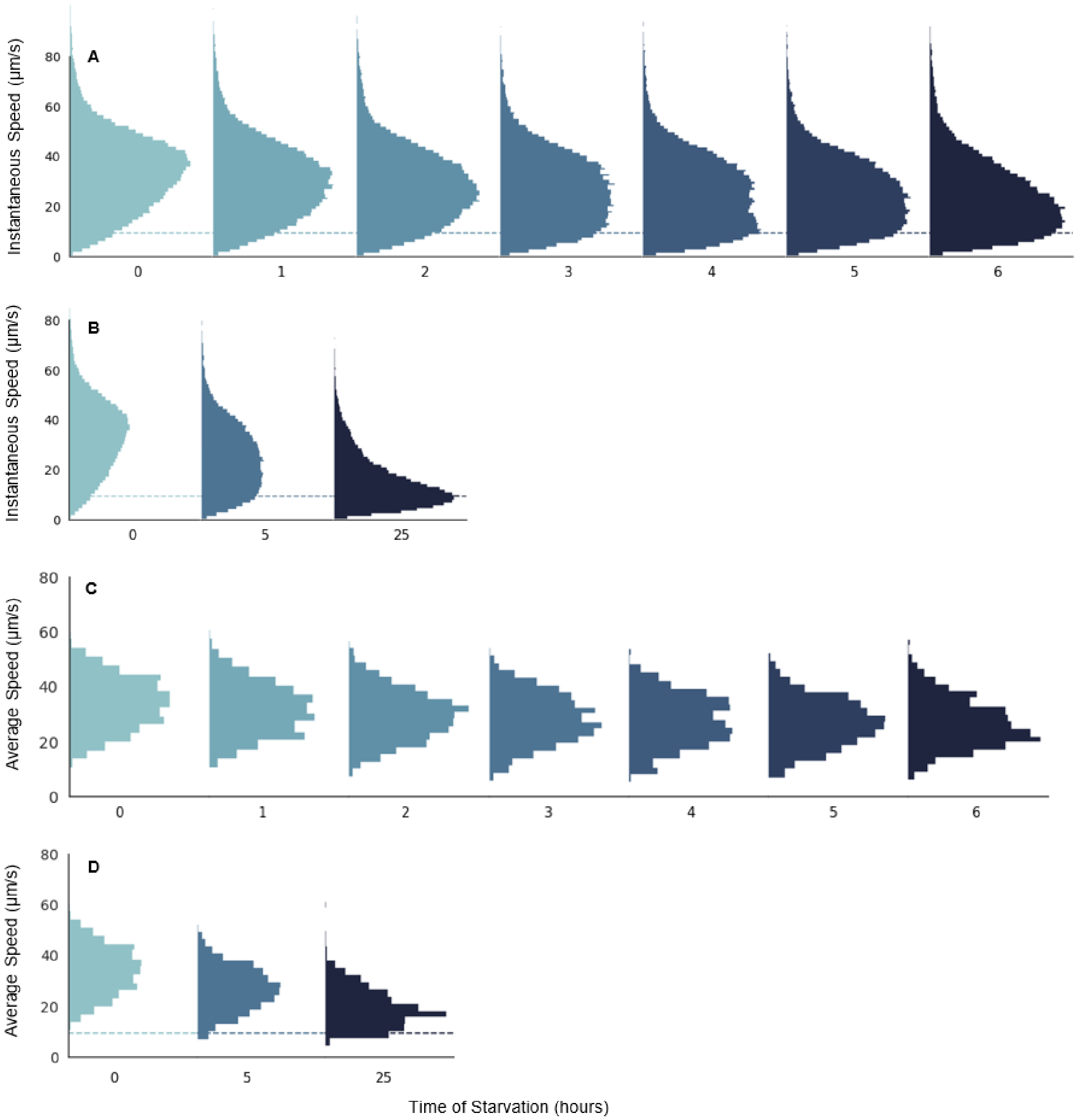


**Fig S3. Under starvation conditions, the instantaneous and average speed distributions of *B. bacteriovorus* shift across time [run 4].** We follow a similar convention to Fig. 1 in the main text. The only relevant difference is the the number of data points contributing to each histogram. For (A) and (B), the number of instantaneous speed observations ranges from 49,264-107,299; for (C) and (D), the number of average speed observations ranges from 500-2,065.

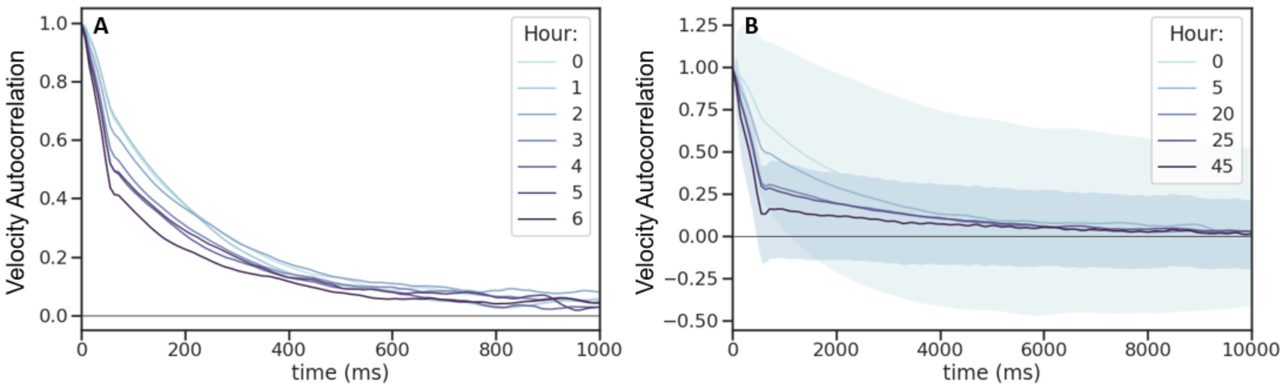


**Fig S4. Under starvation conditions, *B. bacteriovorus'* velocity decorrelates increasingly rapidly as it ages [run 2].** We follow a similar convention to Fig. 3 in the main text.

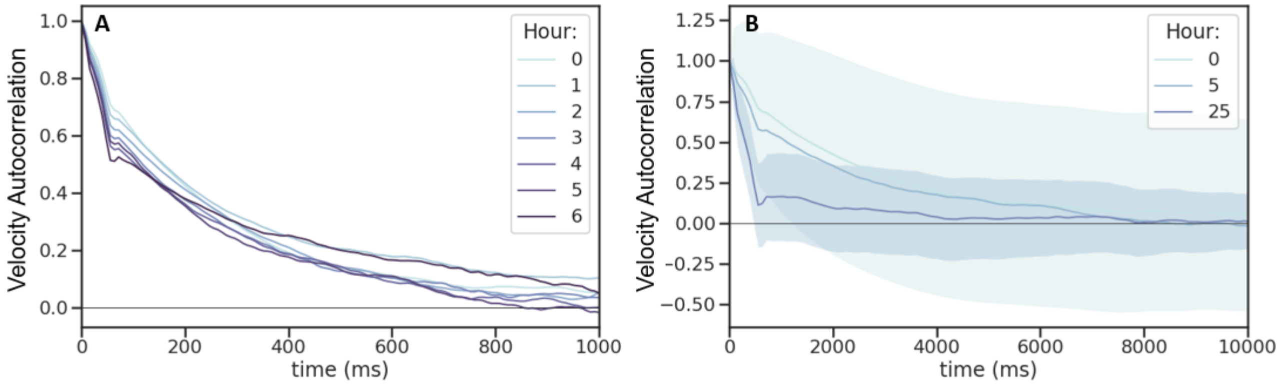


**Fig S5. Under starvation conditions, *B. bacteriovorus'* velocity decorrelates increasingly rapidly as it ages [run 3].** We follow a similar convention to Fig. 3 in the main text.

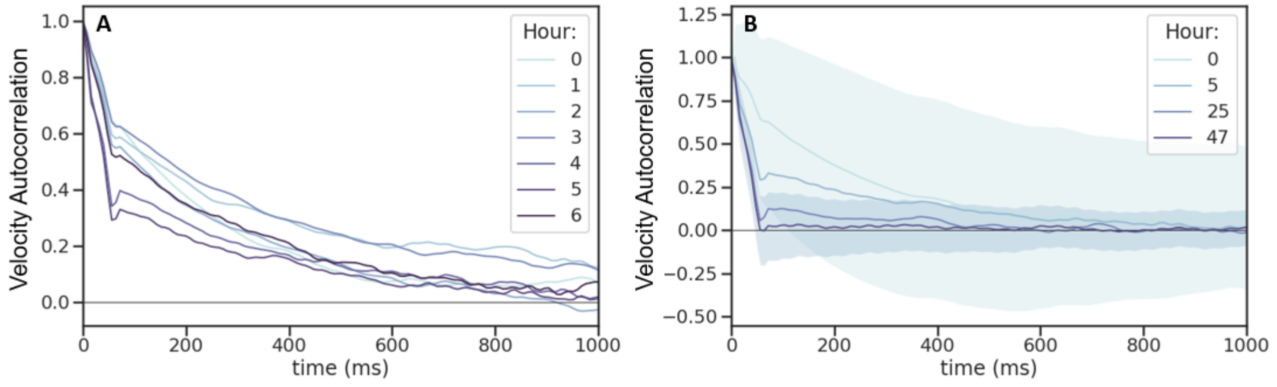


**Fig S6. Under starvation conditions, *B. bacteriovorus'* velocity decorrelates increasingly rapidly as it ages [run 4].** We follow a similar convention to Fig. 3 in the main text.

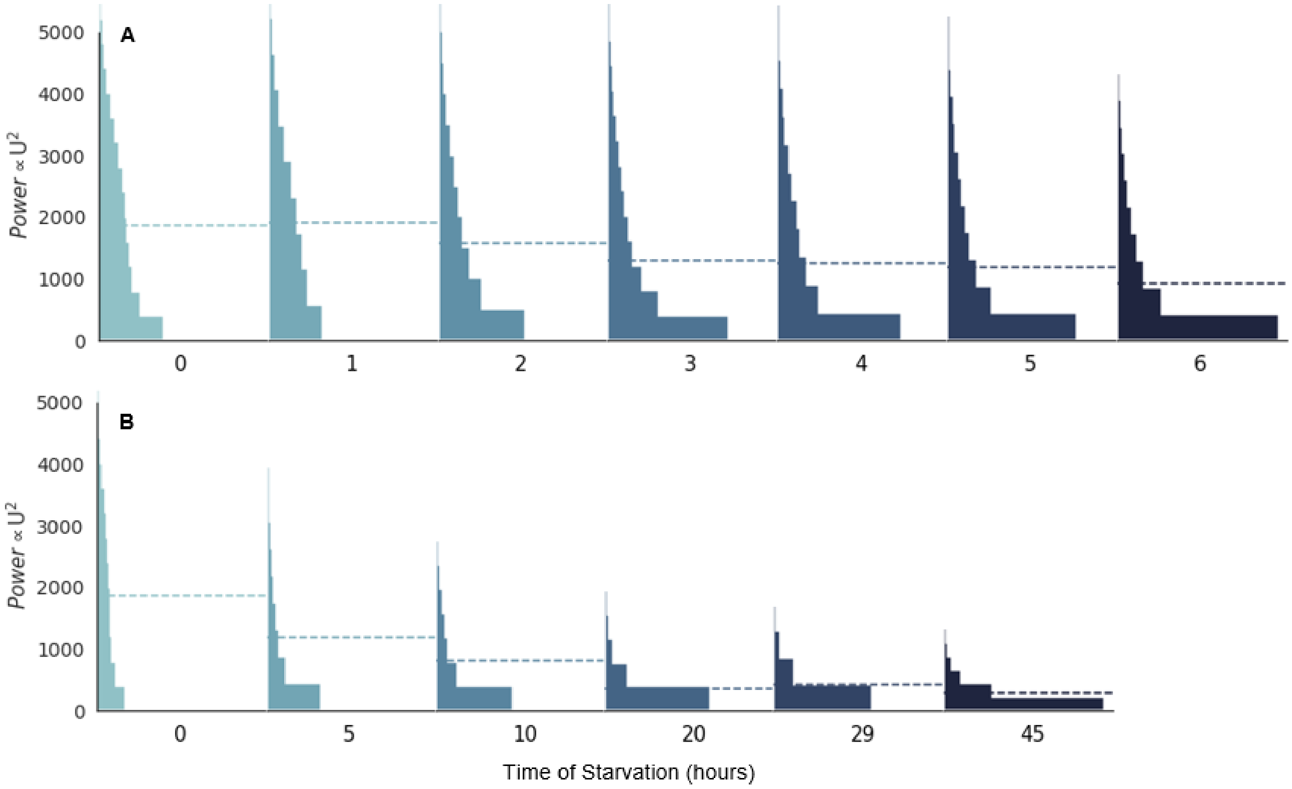


**Fig S7. Under starvation conditions, *B. bacteriovorus* dissipates less power as it ages [run 2].** We follow a similar convention to Fig. 4 in the main text. For (A) and (B), the number of $U^2$ observations ranges from 93,196-172,376.

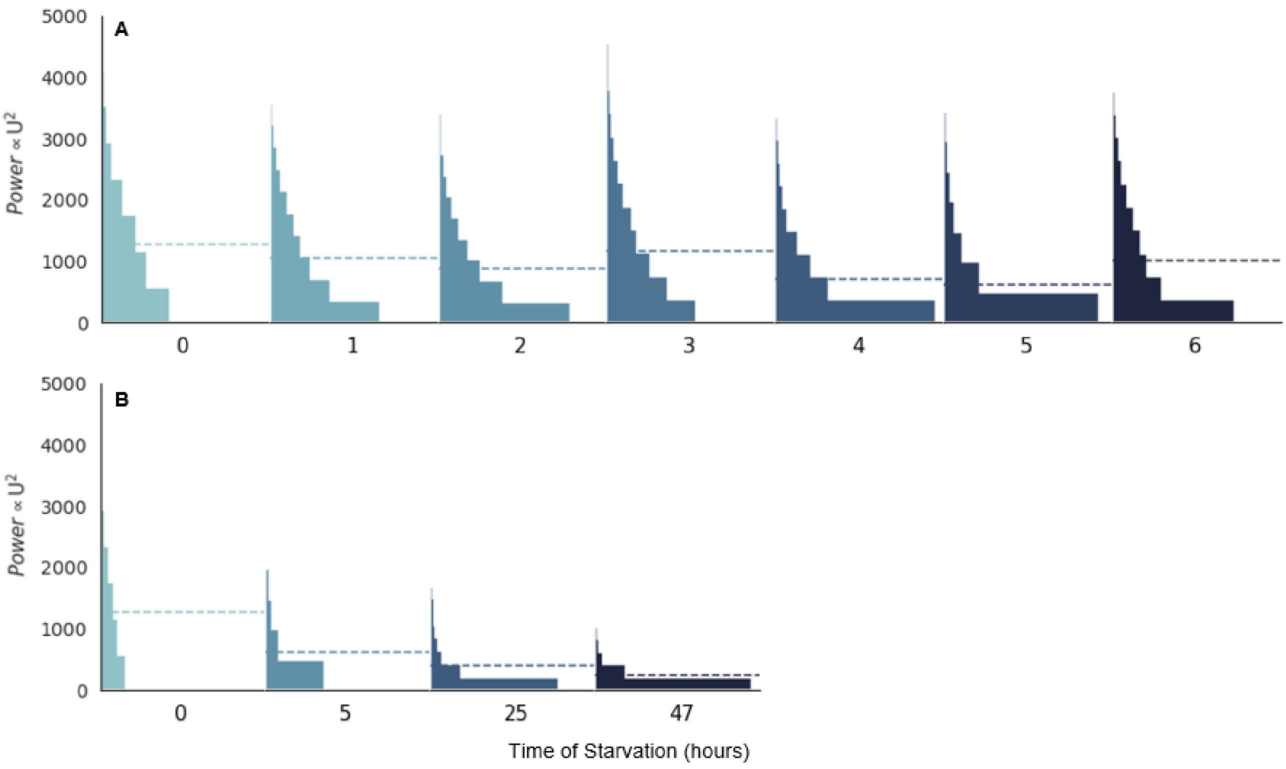


**Fig S8. Under starvation conditions, *B. bacteriovorus* dissipates less power as it ages [run 3].** We follow a similar convention to Fig. 4 in the main text. For (A) and (B), the number of $U^2$ observations ranges from 25,959-60,393.

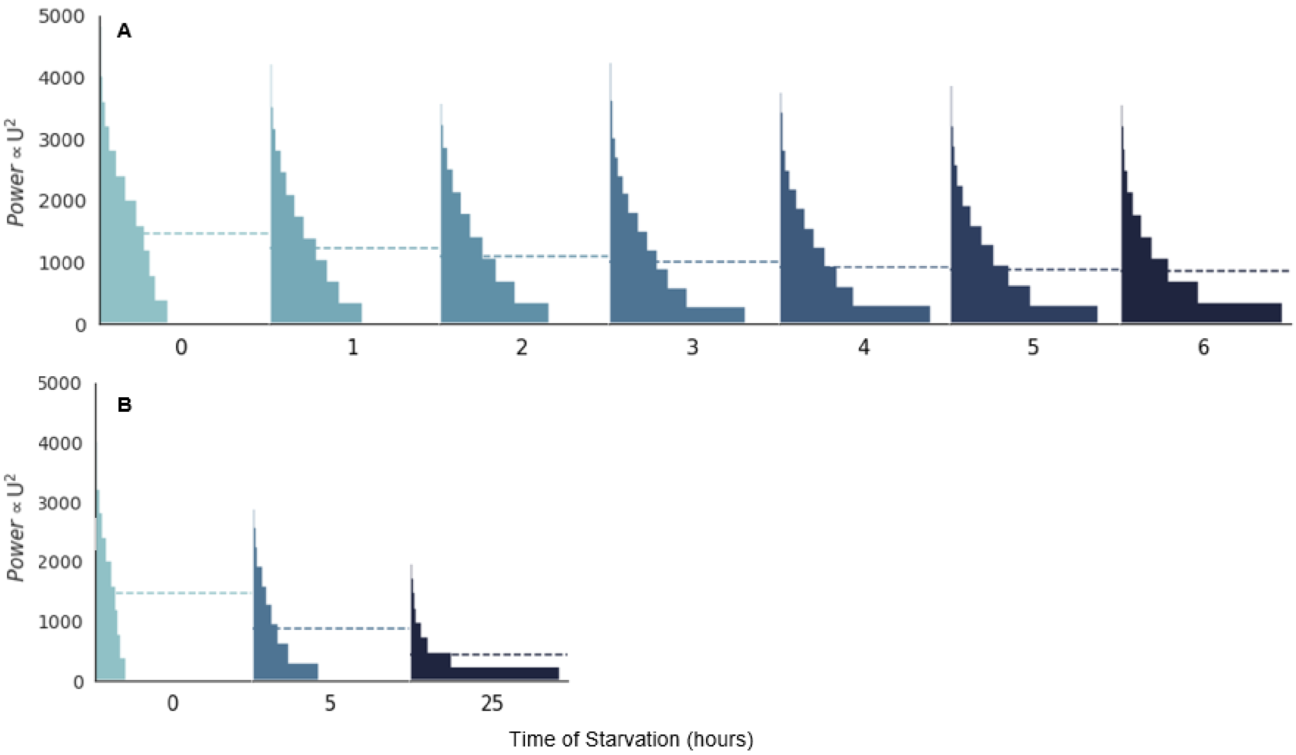


**Fig S9. Under starvation conditions, *B. bacteriovorus* dissipates less power as it ages [run 4].** We follow a similar convention to Fig. 4 in the main text. For (A) and (B), the number of $U^2$ observations ranges from 49,264-107,299.

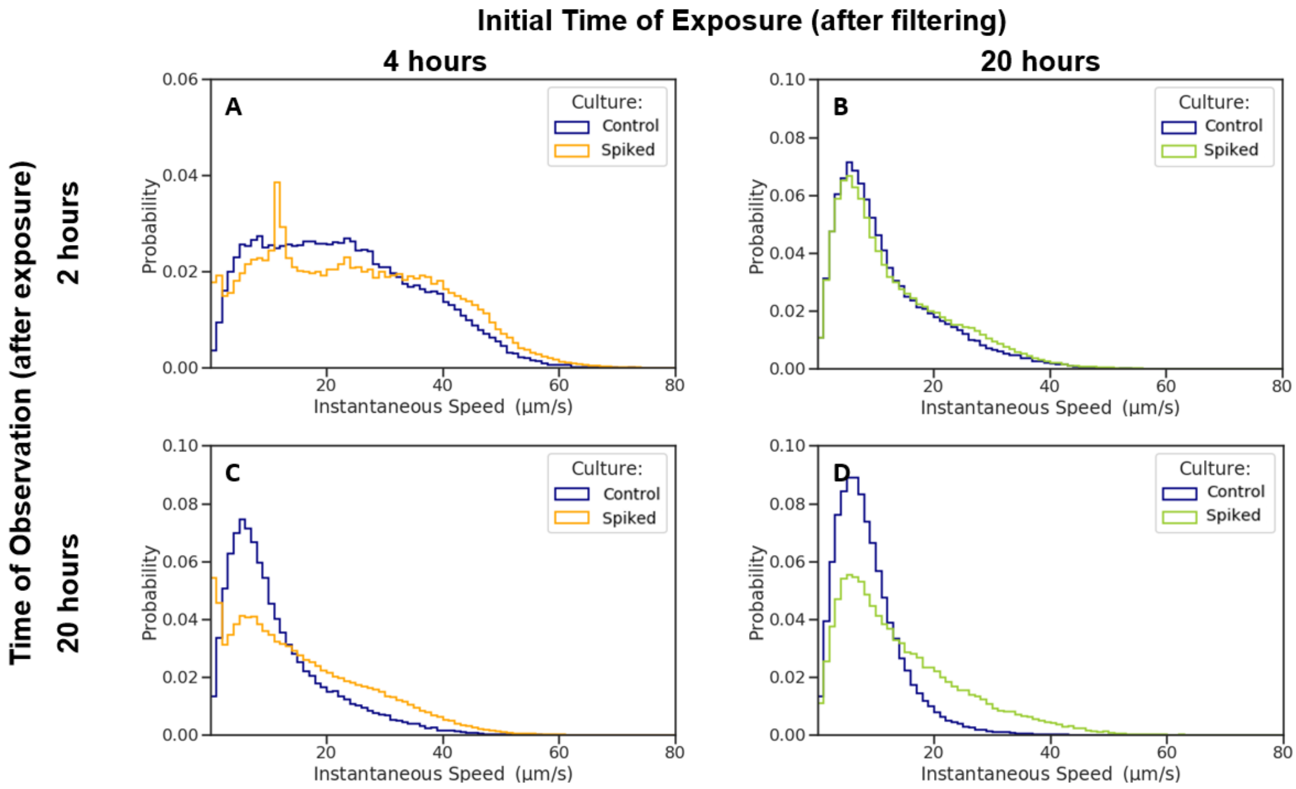


**Fig S10. After addition of LB, *B. bacteriovorus* begins to swim faster even after long starvation times [run 2].** We follow a similar convention to Fig. 5. The number of instantaneous speed observations ranges from 65,795-110,082 for (A), 84,361-174,041 for (B), 80,625-95,618 for (C), and 141,373-96,026 for (D). Distributions were constructed from 100 evenly spaced bins over the range of instantaneous speed measurements.

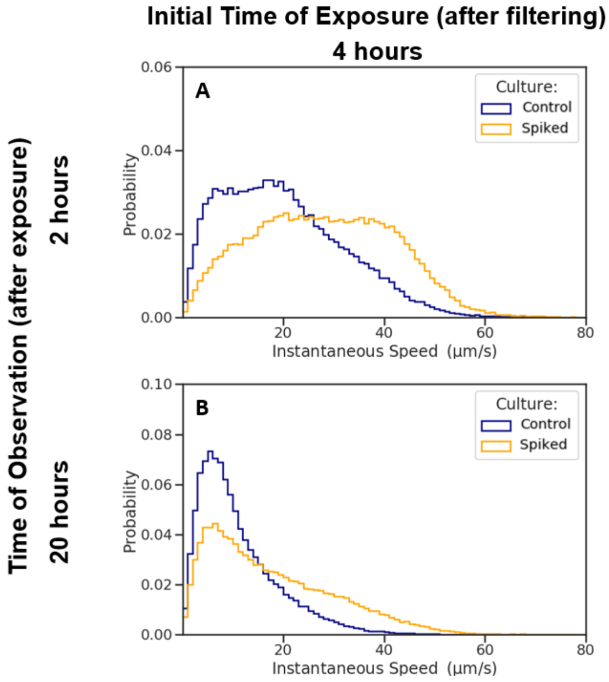


**Fig S11. After addition of LB, *B. bacteriovorus* begins to swim faster even after long starvation times [run 3].** We follow a similar convention to Fig. 5 The number of instantaneous speed observations ranges from 71,290-64,100 for (A) and 68,325-126,841 for (B). Distributions were constructed from 100 evenly spaced bins over the range of instantaneous speed measurements.

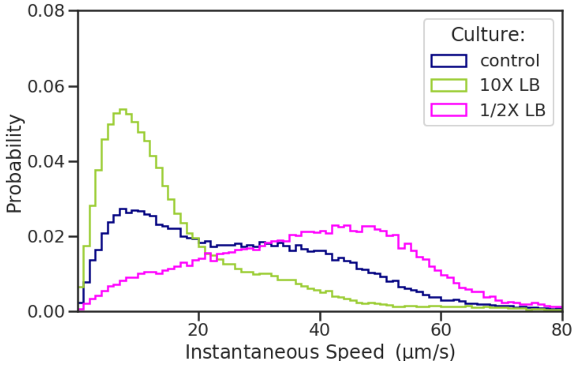


**Fig S12. The speed distributions of spiking experiment is concentration dependent.** A culture starved for four hours and then split into three cultures. One culture remained starving (blue) while the other two were spiked with 10X LB (green) or 1/2X LB concentration (magenta). The culture spiked with the higher concentration resulted in an increase in the slow-speed peak whereas the 1/2X culture exhibited an increase in the higher mode peak. This is likely due to an increase in the osmolarity of the solution with the greater concentration of LB. The number of instantaneous speed observations ranges from 49,390-96,644. The data were divided into 100 evenly spaced intervals.

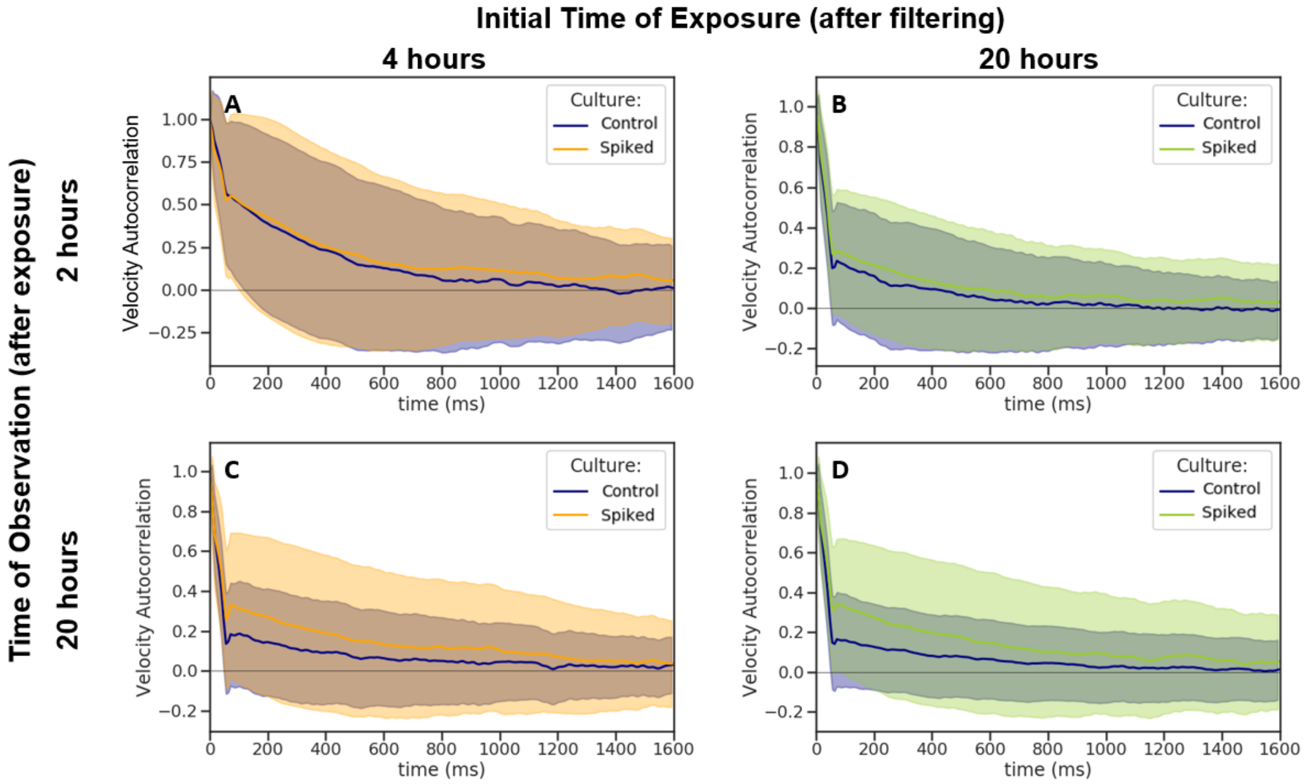


**Fig S13. *B. bacteriovorus* shows a more slowly decaying VACF after the addition of LB [run 2].** We follow a similar convention to Fig. 6.

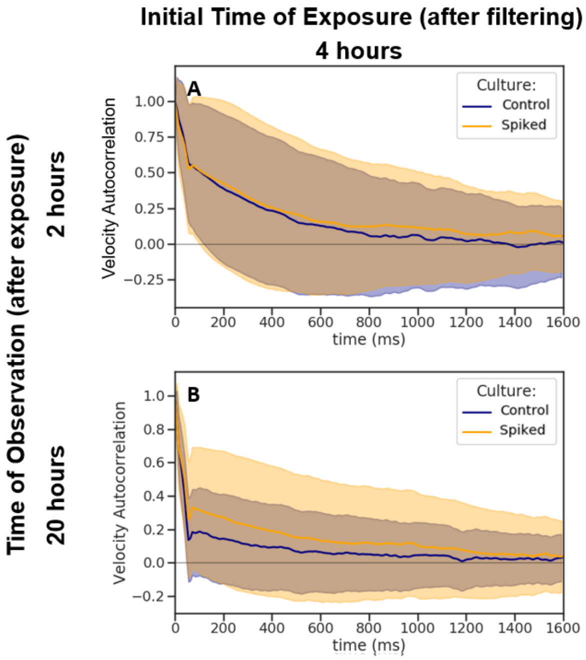


**Fig S14. *B. bacteriovorus* shows a more slowly decaying VACF after the addition of LB [run 3].** We follow a similar convention to Fig. 6.

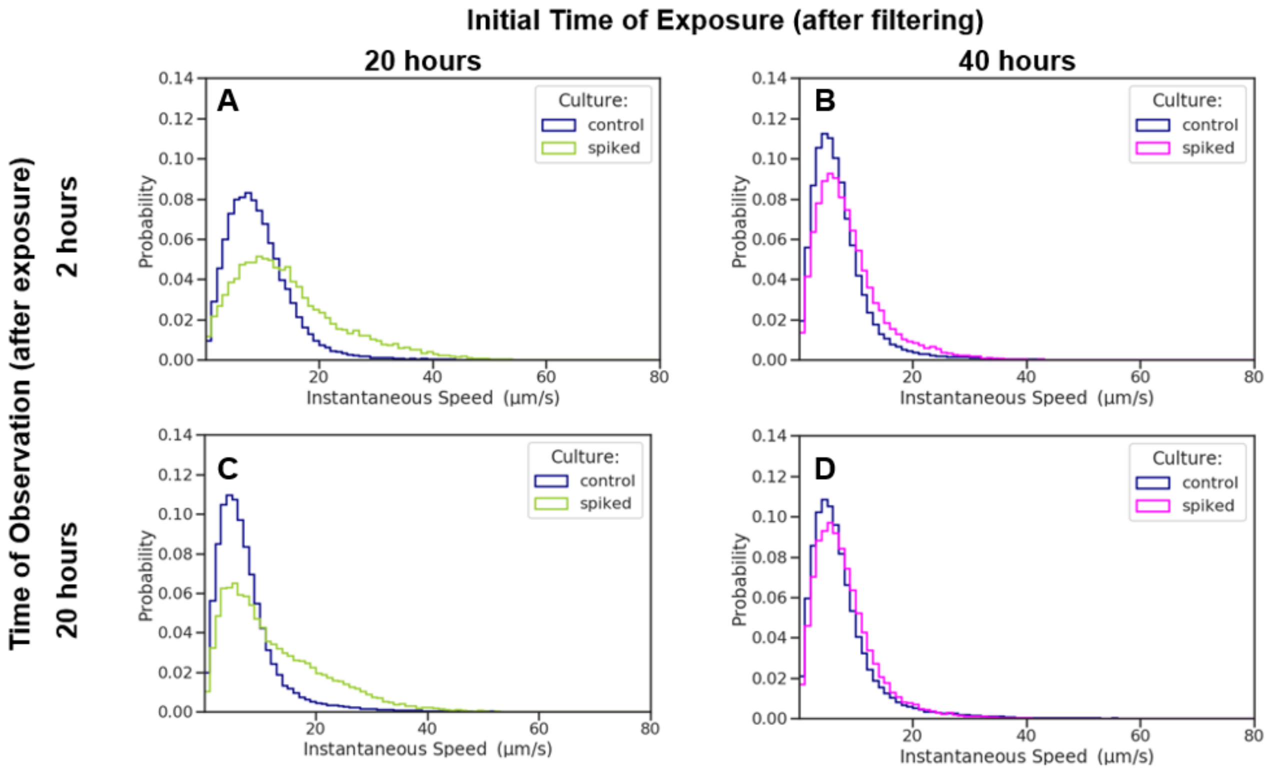


**Fig S15. *B. bacteriovorus* shows no signs of revival after 40 hours of starvation.** We compare the instantaneous speed distributions of our control, starving *B. bacteriovorus* (blue), to that of cultures spiked with LB at Hour 20 (green) and Hour 40 (magenta). Both spiked cultures were observed after 2 hours and 20 hours of exposure. (A) After two hours of exposure, the culture spiked at Hour 20 (green) has faster speeds than the respective control (blue). (B) After two hours of exposure, the culture spiked at Hour 40 has no noticeable change from the control (blue). (C) After 20 hours of exposure, the culture spiked at Hour 20 still has faster instantaneous speeds than the control. (D) After 20 hours of exposure, the culture spiked at Hour 40 has no noticeable change from the control. The number of instantaneous speed observations ranges from 23,472-45,607 for (A), 28,377-79,194 for (B), 46,801-91,407 for (C), and 38,886-84,523 for (D). Distributions were constructed from 100 evenly spaced bins over the range of instantaneous speed measurements.

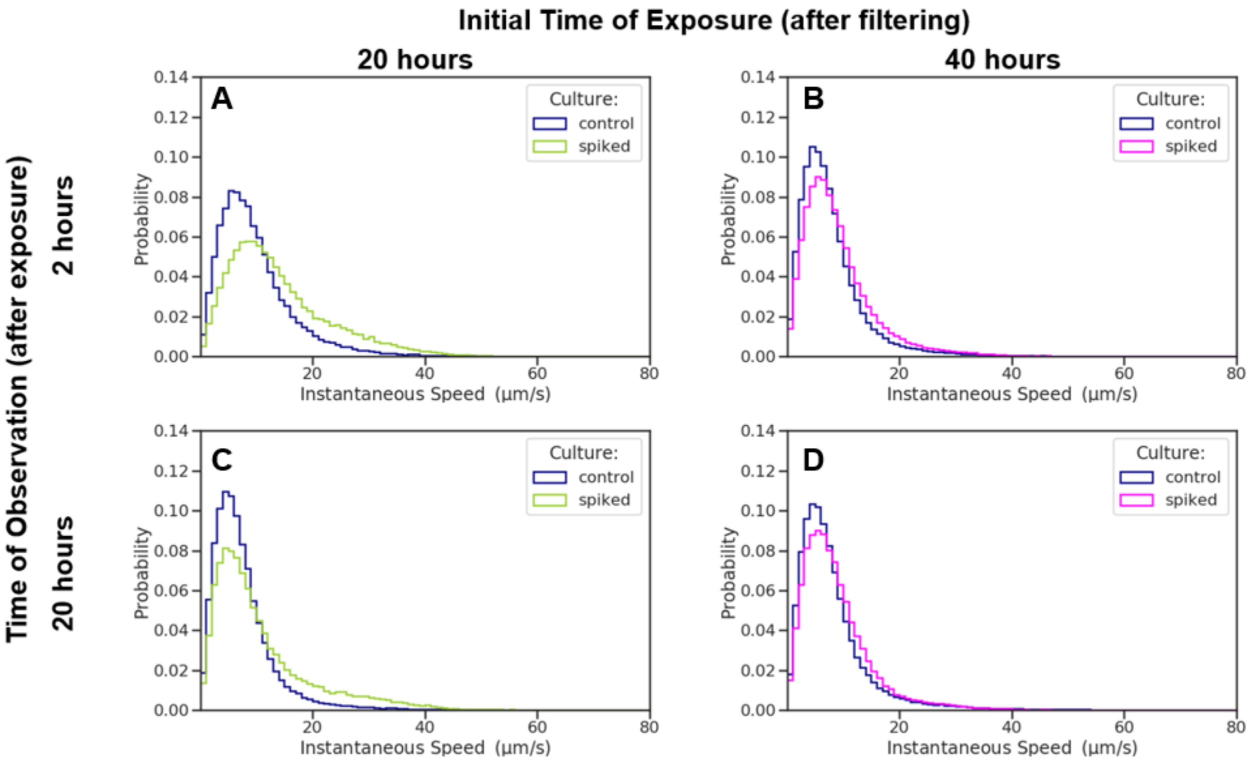


**Fig S16. *B. bacteriovorus* shows no signs of revival after 40 hours of starvation [run 2].** We follow a similar convention to Fig. S15 The number of instantaneous speed observations ranges from 36,346-37,567 for (A), 53,562-114,970 for (B), 72,445-110,723 for (C), and 80,017-188,080 for (D). Distributions were constructed from 100 evenly spaced bins over the range of instantaneous speed measurements.

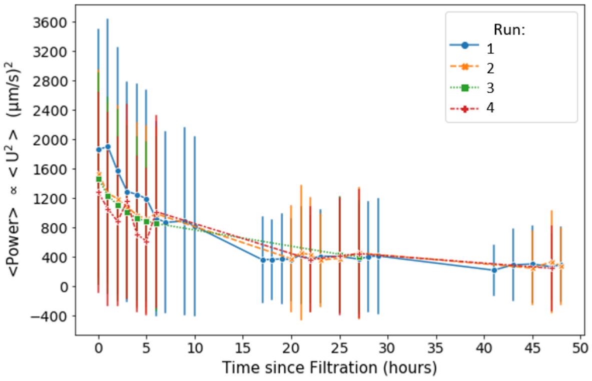


**Fig S17. Different runs result in similar power dissipated by *B. bacteriovorus*.** We follow a similar convention to Fig. 7. Individual runs for the starving culture are plotted. The number of $U^2$ observations ranges from 65,795-151,604 for run 1 (blue), 93,196-172,376 for Run 2 (orange), 25,959-60,393 for Run 3 (green), and 49,264-107,299 for Run 4 (red).

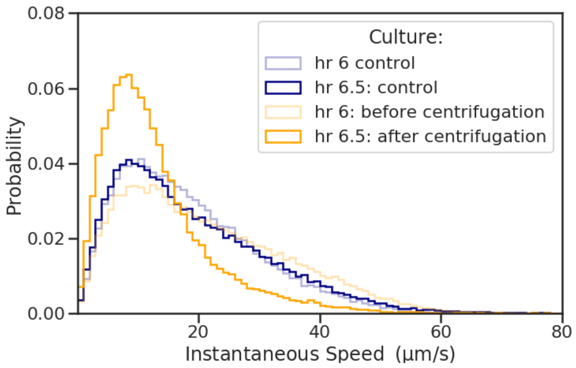


**Fig S18. There is an increase in the population of slow speeds after centrifugation.** A *B. bacteriovorus* culture starved for 6 hours was split as seen in the spiking experiments. One culture was not centrifuged (blue) while the other was centrifuged (orange) as discussed in the memory methods section. There is no noticeable change between the control (blue) at Hour 6 and Hour 6.5. However, there is an increase in the slow speed population after centrifugation (orange).

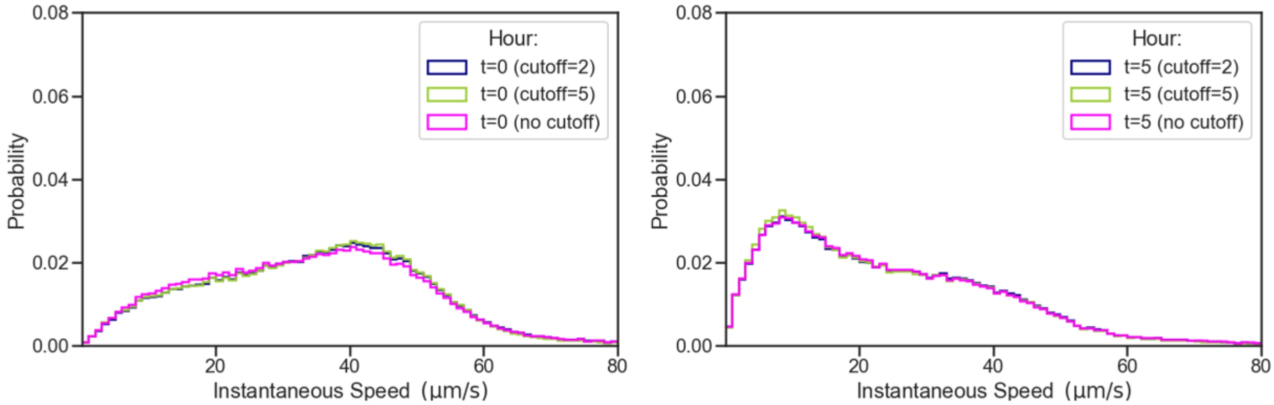


**Fig S19. Varying cutoffs of endpoints of trajectories does not affect the bimodal distribution.** From Table S1, the relative biases of cutting off different endpoints was seen for Hour 0. However, looking at the distributions after removing these frames from each trajectory, there is no noticeable effect on the structure of the bimodal distributions.

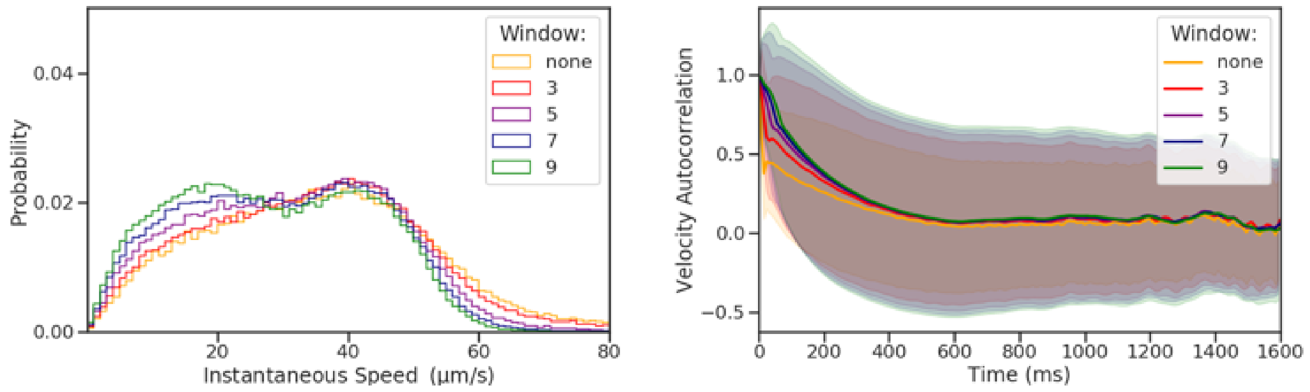


**Fig S20. Using different window averaging has minimal affects on the speed distribution and velocity autocorrelations.** The bimodal distribution for Hour 0 seen in Fig. 1 was observed using different window averaging. Overall a larger window causes the speed distributions to shift slightly to the left, but the overall distribution shape remains the same. The velocity autocorrelation is very similar regardless of the window. The histogram data was divided into 100 evenly spaced bins.

**Table S1. Speeds measured at bounding frames after removal of *n* edge frames.**

| | **Number of frames removed from edge** | | | | | |
|---|---|---|---|---|---|---|
| **Bounding frames** | **0** | **1** | **2** | **3** | **4** | **5** |
| $< 30\,\mu m\,s^{-1}$ | 4303 (55%) | 3310 (43%) | *2885 (37%)* | 2921 (38%) | 2976 (38%) | 3039 (39%) |
| $> 30\,\mu m\,s^{-1}$ | 3464 (45%) | 4457 (57%) | *4882 (63%)* | 4846 (62%) | 4791 (62%) | 4728 (61%) |
| Total | 7767 (100%) | 7767 (100%) | *7767 (100%)* | 7767 (100%) | 7767 (100%) | 7767 (100%) |

Speeds measured at the beginning or end of a trajectory are biased towards lower speeds because bacteria moving into or out of focus are more likely to have a $z$-component to their velocity while motion occurs in the $xy$-plane. To determine the degree of bias, between 0 and 5 *edge frames* were trimmed from the beginning and end of all trajectories and the speeds re-measured for the three bounding frames (at the beginning and end of each trajectory). The number (and percentage) of samples above and below $30\,\mu m\,s^{-1}$ are counted after trimming a given number of edge frames. Removing roughly two edge frames (emphasized above) is sufficient to compensate for the slow speed bias at the ends of trajectories; the percentage of slow speed samples settles around 38%.

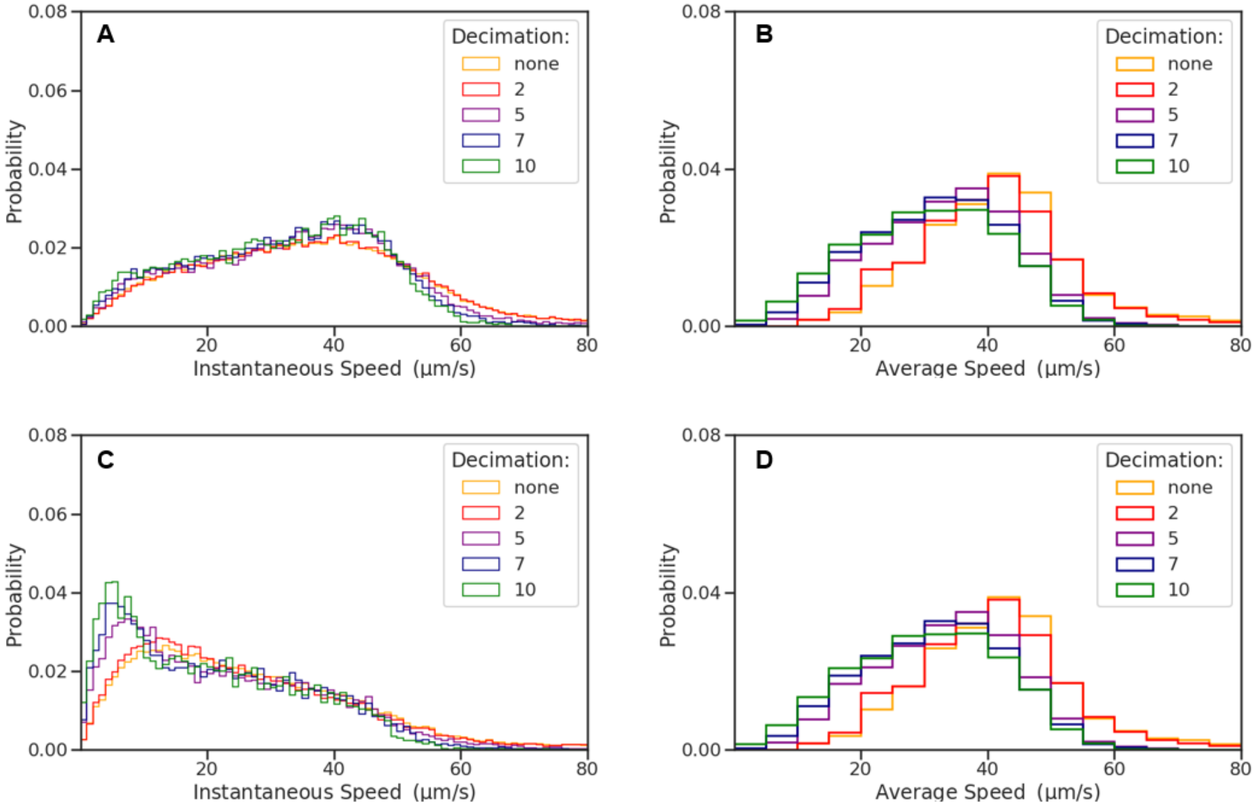


**Fig S21. Decimating the data has minor effects in the speed distributions.** To further test the bimodal behavior seen at early hours of starvation, the data was decimated. For example, for a decimation of 5, every fifth data point of a trajectory was used, thus also decreasing the number of samples. (A) Decimating the data does not show a noticeable effect even as much as taking only every tenth instantaneous speed of a bacterial trajectory at Hour 0. (B) A slight shift to a slower average speed is seen when utilizing less data points, but the overall shape of the distribution does not vary greatly at Hour 0. (C) Decimating the data at Hour 5 has an increase in the height of the slow peak when taking every fifth to tenth data points. (D) The average speed distributions also show a shift to the left at Hour 5 with larger decimations. As the number of data points is less for Hour 5 compared with Hour 0, this may be a cause for further effects of using larger decimations (less data points). All data in the instantaneous speed and average speed histograms was divided into 100 and 30 evenly spaced bins, respectively.

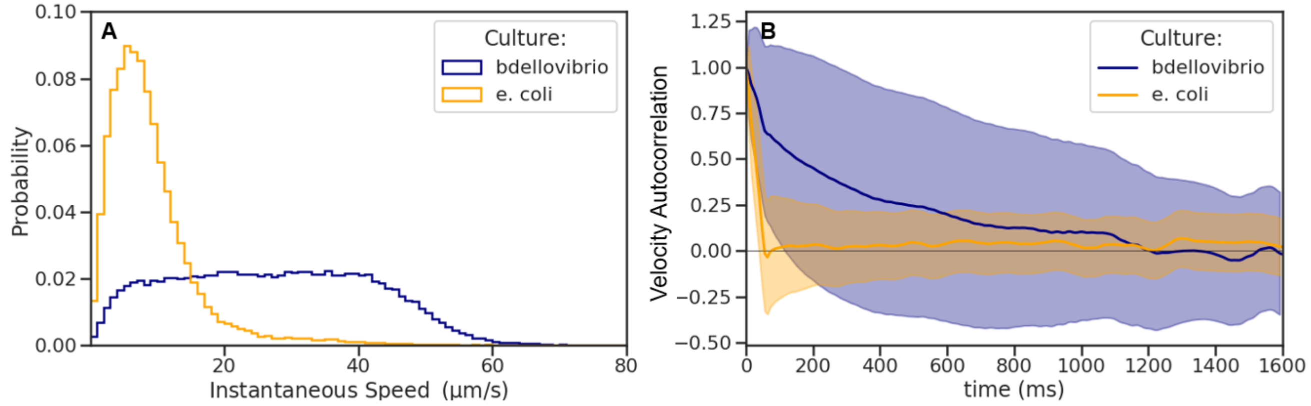


**Fig S22. *E. coli* speed distributions and VACFs greatly differ from *B. bacteriovorus* at early exposure.** *E. coli* strain OP50 was starved for one hour and then spiked with LB. *B. bacteriovorus* was starved for four hours and observed one hour thereafter. (A) Although there is a slight variation in the starvation times, *E. coli* does not show any bimodal behavior in the instantaneous speed distribution. (B). The velocity of *E. coli* also becomes decorrelated much faster than *B. bacteriovorus*, indicating that while the predator is swimming, the prey is diffusing. Therefore, if there were leftover *E. coli* in the solution, a much greater increase in the slow peak would be observed. As we do not seen this upon exposure to LB with our *B. bacteriovorus* spiking runs, we can further conclude that *E. coli* is not present nor contributing to the distributions. The data was divided into 100 evenly spaced bins.

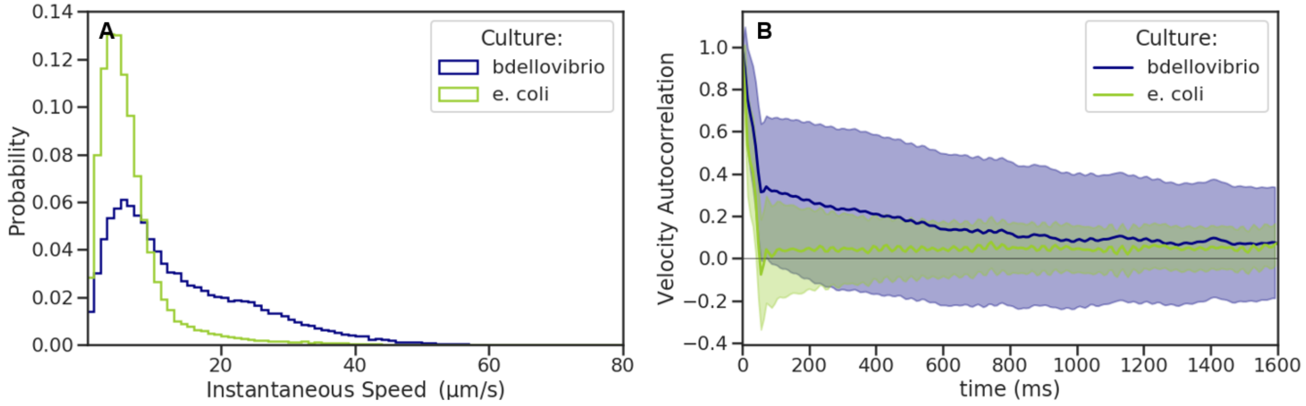


**Fig S23. *E. coli* speed distributions and VACFs greatly differ from *B. bacteriovorus* at late exposure.** Separate cultures of *E. coli* strain OP50 and *B. bacteriovorus* were starved for twenty hours and then spiked with LB. Both cultures were observed three hours after exposure. (A) Again, *E. coli* does not show any bimodal behavior in the instantaneous speed distribution. (B). The velocity of *E. coli* becomes decorrelated much faster than *B. bacteriovorus*. Therefore, even after longer exposure to LB, OP50 is still not motile as seen with the predator. The data in the histogram were divided into 100 evenly spaced intervals.
